# Core circadian clock genes control molecular and behavioral circatidal rhythms in *Parhyale hawaiensis*

**DOI:** 10.64898/2026.02.27.708297

**Authors:** Victoria Louis, Zachary Bellido, Adam Helfenbein, Joshua J. C. Rosenthal, Patrick Emery

## Abstract

Marine organisms exhibit 12.4-hour rhythms of gene expression, physiology and behavior synchronized by tidal cues. The mechanism underlying these circatidal rhythms, and its overlap with the circadian clockwork, has remained elusive. However, recent studies showed that the core circadian gene BMAL1 sustains circatidal behavior in crustaceans. Therefore, we mutagenized the other three core circadian clock genes (*PhCry2, PhPer* and *PhClk*) in *P. hawaiensis*, a marine amphipod. We found that they are necessary for both circadian and circatidal behaviors. Moreover, all four core circadian genes are critical for 24-h oscillations of mRNA levels in circadian brain neurons and 12.4-h mRNA rhythms in circatidal neurons. Unexpectedly, the mutants indicate that PhCLK represses *PhPer* expression independently of PhBMAL1 specifically in circatidal neurons. Our study thus reveals that circadian and circatidal clocks share four core molecular components, but their transcriptional wiring differs.

## Introduction

Organisms adjust their metabolism, physiology and behavior to predictable cycles in their surrounding environment using biological clocks ^1–4^. Among them, circadian clocks are by far the best understood. They are entrained by the alternance of day and night and free-run with a ∼24-hour (h) period when held in constant environmental conditions. The circadian clock is highly conserved across animals^1,5^. Its core is a negative transcriptional feedback loop comprised of the activators CLOCK (CLK) and BMAL1 (called CYCLE in *Drosophila*) that heterodimerize to bind to E-boxes and activate the transcription of numerous genes, including those encoding their own repressors PERIOD (PER, CRYPTOCRHOME (CRY) and/or TIMELESS (TIM), depending on the species. Other CLK/BMAL1 targets include transcription factors that form interlocked loops that modulate and strengthen the core loop ^5–11^.

In marine environments, especially in the intertidal zone, organisms are subjected to drastic environmental changes caused by tides. In response, besides circadian rhythms, they also exhibit ∼12.4-h rhythms that synchronize to tidal cues ^12,13^. The molecular mechanism behind these circatidal rhythms remain poorly characterized. Early behavioral studies of different intertidal organisms led to three main hypotheses. The first posited that circadian and circatidal rhythms are generated by distinct clocks ^14^. The second proposed that circadian and circatidal rhythms are produced by a single clock, which can run with a period of ∼24 h or ∼12.4 h, depending on the rhythmic cues an animal is exposed to ^15^. The last hypothesis suggested the existence of two circalunidian (∼24.8-h) clocks running in antiphase, thus generating circatidal (∼12.4-h) outputs ^16^. This third hypothesis could involve essentially the same mechanism as the circadian clock, with a minor adjustment of the oscillator’s period (from 24 to 24.8h). A similar minor period adjustment could also permit the circadian clock mechanism to generate 12.4-h period rhythms by exploiting its two anti-phasic interlocked transcriptional feedback loops, or by driving neuronal activity in opposite phase in two different neuronal groups, as observed in *Drosophila* in Morning and Evening activity promoting neurons ^17,18^.

Several studies have thus tested whether circadian clock genes are required for circatidal rhythms. Knock downs of core circadian clock genes via RNA interference (RNAi) were performed in the mangrove cricket, *Apteronemobius asahinai*, and in the marine isopod *Eurydice pulchra*. Results suggested that the core circadian genes Clk, Per and Cry2 are not required to sustain circatidal outputs ^19–22^. However, pharmacological inhibition of Casein Kinase Iε, which phosphorylates PER proteins and thus regulates circadian rhythms in flies and mammals ^23^, lengthened circatidal behavioral rhythms, suggesting that this kinase is shared between the circadian and circatidal clocks ^21^. More recent studies further point towards a mechanistic overlap between the two clocks. Indeed, CRISPR-Cas9 mediated knock out of *PhBmal1* in the amphipod *Parhyale hawaiensis* eliminated circatidal behavior ^24^. Moreover, RNAi directed at Bmal1 reduced circatidal rhythmicity in *E. pulchra*. Recently, distinct circadian and circatidal clocks cells were identified in the brain of *P. hawaiensis* and *E. pulchra* ^25^. Results were particularly striking in *P. hawaiensis*, as circatidal (12.4-h) rhythms of expression for both *PhPer* and *PhCry2* mRNAs were observed in four out of the ∼60 cells expressing core circadian genes. Moreover, these cells synchronize with a tidal cue (i.e. vibration), but not to the light/dark cycle. By contrast, circadian cells displayed 24-h period rhythms of *PhPer* and *PhCry2* mRNAs that synchronized to the LD cycle, but not to vibration. These data suggest that *PhPer* and *PhCry2* may contribute to circatidal rhythms in *P. hawaiensis*.

In the present study, we used CRISPR-Cas9 genome editing to test the degree of mechanistic overlap between the circadian and the circatidal clocks in *P. hawaiensis* by generating knock-out lines for *PhPer, PhClk* and *PhCry2* to complement the existing *PhBmal1* knock-out animals. Interestingly, we found that all four core clock genes are necessary to sustain circatidal rhythms, indicating considerable mechanistic overlap between the two clock mechanisms. Nonetheless, our results also suggest that the transcriptional wiring between the four core clock genes differ between circadian and circatidal cells.

## Results

### Loss of *PhCry2* disrupts circadian behavior

Based on genomic studies, the circadian clock of *P. hawaiensis* is predicted to comprise homologs of CLK, BMAL1, PER and repressor-type cryptochromes ^26^. The genome does not appear to encode a photoreceptive CRY or a TIM homolog. We previously demonstrated that BMAL1 is required for both circadian and circatidal behavioral rhythms. We therefore decided to determine whether the circadian and circatidal clocks share additional common building blocks.

We first targeted the *PhCry2* gene. Two knock-out lines were generated using CRISPR-Cas9 and a set of two guide RNAs (gRNA) targeting the photolyase domain (Figure 1A), which is required for cryptochromes’ repressor function ^27^. The first mutant line (*PhCry2*-mutation 1) harbors two deletions of 18 and 20 base pairs (bp), respectively, which cause a premature stop codon that truncates the PhCRY2 protein from 929 amino acids (AA) to 422 AA. The second line (*PhCry2*-mutation 2) harbors a single deletion of 16 bp, also causing a premature termination codon that truncates PhCRY2 to 428 amino acids.

**Figure 1:**
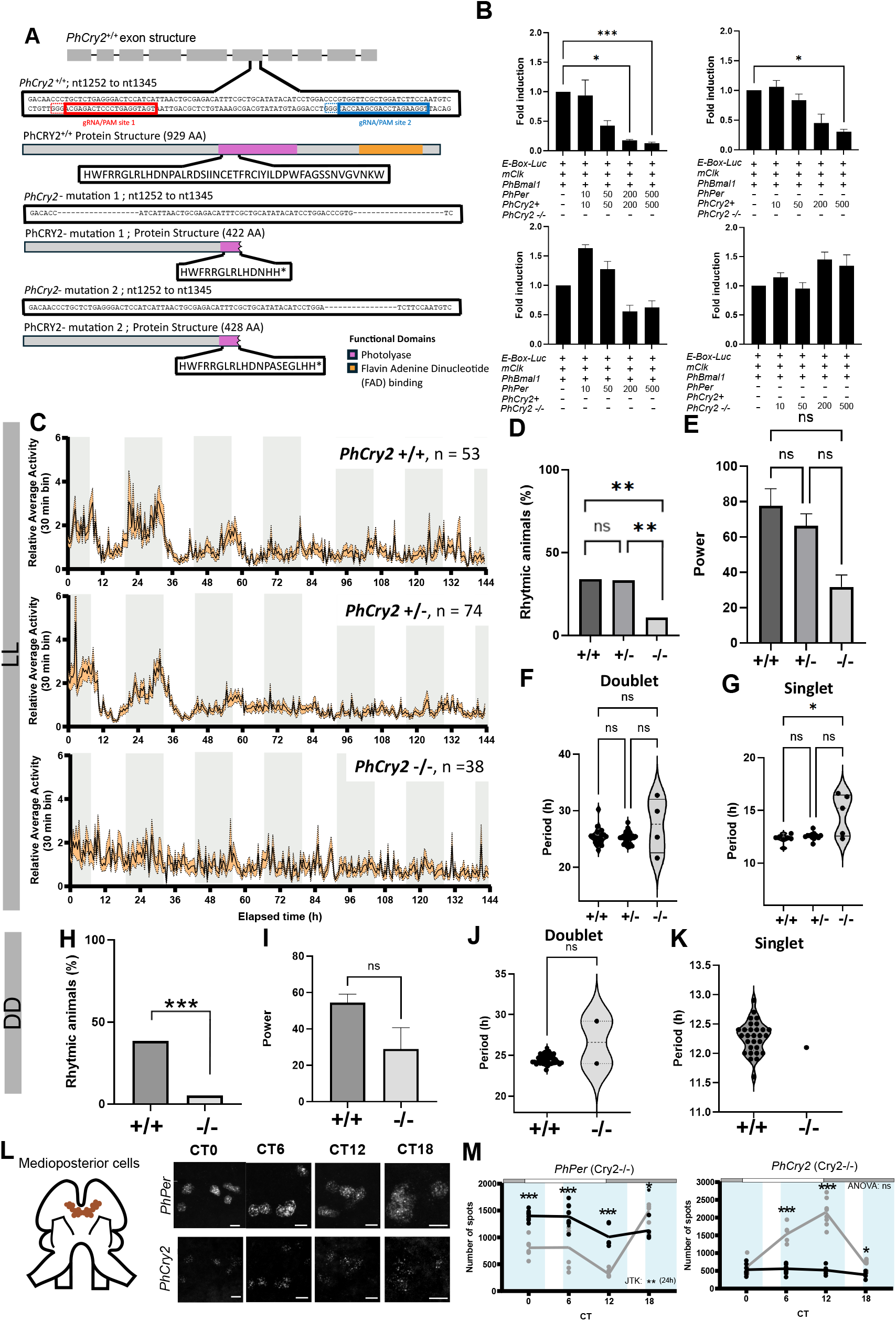
PhCRY2 is required to maintain circadian rhythms in *P. hawaiensis*. A) Schematics of *PhCry2* mutagenesis. Blue and red boxes indicate the gene sequences targeted by the gRNAs used for CRISPR/Cas9 mutagenesis. Two knock-out mutant lines with premature stop codons were obtained. The resulting mutant PhCRY2 proteins are very similar, with slightly different C-terminal tails (see boxed amino acid sequences). Both are missing most of the photolyase domain (pink) and the Flavin Adenine Dinucleotide (FAD) binding domain (orange). Thus, behavior data from both strains were combined on this figure and on figure 2. B) Reconstruction of the circadian clock mechanism in HEK-293T cells with luciferase-based transcriptional assay. Co-transfection of increasing concentrations (in ng) of *PhPer* and *PhCry2* increase the repression of *mClk*:*PhBmal1*-mediated transcription (Kruskal-Wallis test, ***p<0.001; subsequent Dunn’s multiple comparison, 0 ng vs 200 ng, *p=0.023, 0 ng vs 500 ng, ***p=0.001; N=5). The transfection of *PhCry2* alone can repress as well (Kruskal-Wallis test, **p=0.009; subsequent Dunn’s multiple comparison, 0 ng vs 500 ng, *p=0.023; N=4). *PhPer* alone does not repress (Kruskal-Wallis test, p=0.002; subsequent Dunn’s multiple comparison p = ns; N=4). The mutated version of *PhCry2* (−/−) does not repress transcription (Kruskal-Wallis test, *p=0.018; subsequent Dunn’s multiple comparison shows no significant differences compared to control; N=3). C) Average relative swimming activity of wild-type (*PhCry2*+/+), heterozygous (*PhCry2*+/-) and homozygous (*PhCry2*−/−) animals recorded in constant light (LL) after entrainment to a LD cycle. Gray shading represents the subjective night. Standard error to the mean (SEM) is represented with the orange shading. Vertical swimming^24^ traces are shown in all main figures. See supplemental figures for roaming behavior^24^. D) Percentage of rhythmic animals in LL (Chi-square test, **p = 0.01; subsequent pairwise Chi-square test, +/+ vs +/-, p = 0.94, +/+ vs −/−, **p = 0.004 and +/- vs −/−, **p = 0.005). E) Rhythm power averages under LL (One-way ANOVA, p = 0.0677). F) Doublet period averages under LL (Kruskal-Wallis test, p = 0.670). G) Singlet period averages under LL (Kruskal-Wallis test, *p = 0.031; subsequent Dunn’s multiple comparison, +/+ vs +/-, p = 0.421, +/+ vs −/−, *p = 0.026 and +/- vs −/−, p = 0.345). H) Percentage of rhythmic animals under DD after LD entrainment (Chi-square test, ***p < 0.001; subsequent pairwise Chi-square test, ***p < 0.001) Note that the same control animals were used in Figures 3 and 5. Therefore statistical analysis for the DD datasets were performed with multiple-comparisons tests that included all 4 genotypes (Wild-type, *PhCry2, PhPer* and *PhClk* mutants). I) Rhythm power averages under DD (Mann-Whitney test, p = 0.231). J) Doublet period averages under DD (Mann-Whitney test, p = 0.730). K) Singlet period averages under DD. L) HCR-FISH of *PhPer* and *PhCry2* mRNAs in circadian medioposterior clock neurons exposed to constant conditions after entrainment to 10.3:2.1 tidal and 12:12 LD cycles in *PhCry2* −/− animals (*PhCry2* – mutation 1). A z-stack for each mRNA visualized in the left hemisphere is shown across circadian time (CT). Scale bars represent 10µm. M) Quantification of the number of HCR-FISH spots per hemisphere across CT in *PhCry2* −/− animals. Each dot represents one hemisphere of one individual and the solid line the mean. Grey shading represents the subjective dark phase and blue shading the subjective high tide. Grey trace represents the WT and black trace the mutant. After testing the effect of time on gene expression (*PhPer*, Kruskal-Wallis, **p = 0.0072, *PhCry2*, One-way ANOVA, p = 0.205), rhythm was tested using the JTK-cycle algorithm for 24-h periodicity for *PhPer* (See Table S2 for exact p-value). Statistical differences between genotypes are indicated with * signs (Tukey’s multiple comparison, See Table S3 for exact p-value). Significance thresholds: *p<0.05, **p<0.01 and ***p<0.001 See also Figures S1, S2 and Table S2, S3 and S6

To verify that the truncation affects PhCRY2 activity, we turned to a transcriptional assay in mammalian HEK-293T cells (Figure 1B). We previously showed that PhBMAL1 can activate E-bx mediated transcription with the help of mCLK^24^ (there is still uncertainty on the exact sequence of PhCLK, see below). We thus tested if PhPER and PhCRY2 can repress PhBMAL1/mCLK, as observed in other animals ^21,28–32^. Indeed, we found that expression of both PhPER and PhCRY2 repressed PhBMAL1/mCLK activity in a dose-dependent manner. Repression was also observed with PhCRY2 alone, but it was not as efficient as with PhPER (Two-way ANOVA, *p=0.01). In addition, PhPER did not repress on its own. Thus, as in other animals, CRY2 appears to be the primary repressor, while PER plays a modulatory function ^29,30,32,43^. No repression was observed with expression of the truncated PhCRY2 −/−, which is thus severely defective.

To assess whether the loss of PhCRY2 disrupts the circadian clock as expected, we measured the swimming activity of mutants and control animals under constant conditions after entrainment to a 12:12 light-dark (LD) cycle. We prioritized observing free running behavior under constant light (LL), as we previously observed that circadian behavioral rhythms are more robust under these conditions than under constant darkness (DD) ^24^. This was also the case in the present study (Figure 1C and S1A), although rhythms were weaker than previously reported under both conditions. As expected, under LL, average activity traces of wild-type (WT) animals (*PhCry2* +/+) showed lower activity during the subjective light phase and higher activity during the subjective dark phase (Figure 1C and S2A). The heterozygous animals (*PhCry2* +/-) exhibited a similar pattern. In contrast, the average swimming activity of the homozygous animals (*PhCry2* −/−) showed no obvious rhythmicity. This was reflected in the proportion of rhythmic animals (Figure 1D) and in the rhythm power (amplitude) which was quite weak in the rare rhythmic knockout animals (Figure 1E). We next determined the period of behavioral rhythms using periodogram analysis. After LD-only entrainment, most rhythmic animals exhibit statistically significant rhythm periodicities in the circadian (∼24-h) range (“doublet”), but harmonics in the ∼12-h range can also be observed (“singlets”) ^24^. The average circadian doublet period of WT animals was around 25.5 h as previously described under LL, since continuous light exposure lengthens circadian rhythms in *P. hawaiensis* ^24^. While the average circadian period did not statistically differ between wild-type, heterozygous and the rare rhythmic homozygous mutants, individual periodicities were not clustered around 25.5 h in mutant animals, as in wild-type and heterozygous animals (Figure 1F). Actually, out of the four rhythmic *PhCry2* −/−, only one had a period within the expected range. The singlet period of the homozygous was also very variable, and statistically significantly longer than WT (Figure 1G).

To further establish that PhCRY2 controls circadian rhythms, we tested LD-only entrained animals in constant darkness (DD). Again, circadian behavior was severely disrupted (Figure S1). Only two knockout animals out of 32 were rhythmic and their period was not in the expected range (Figure 1H to 1K). Therefore, our results show that in the absence of functional PhCRY2, circadian behavior is severely compromised. As previously observed with *PhBmal1*−/− mutant animals, residual rhythms of low amplitudes and poorly defined period emerge in a few animals when the circadian clock is disrupted ^24^.

### Loss of *PhCry2* disrupts circadian molecular rhythms

In *Drosophila*, ca. 240 neurons rhythmically express circadian clock genes^33^. These neurons drive different 24-h metabolic and behavioral rhythms, including the timing of the sleep/wake cycle. In addition, numerous glial cells express core clock genes ^34–36^. In the Parhyale brain, only ∼ 60 cells detectably express *PhPer* and *PhCry2* ^25^. To assess the effect of the *PhCry2* knock-out on the circadian clock at the molecular level, we turned to HCR-FISH to monitor *PhPer* and *PhCry2* mRNA rhythms in putative circadian neurons, the medioposterior (MP) cells ^25^. These cells were found to entrain to the LD cycle, showing 24-h mRNA abundance rhythms after entrainment to both a 12:12 LD cycle and a 12.4-h vibration cycle that mimics tides ^25^. Using probes to the pan-neuronal marker elav, we established that most MP cells indeed are neurons, although 1/10 cell per hemisphere appeared elav-negative (Figure S3A). Since we use immersion cycles instead of vibration for tidal entrainment, we first verified that the MP neurons exhibit circadian mRNA rhythms in our hands. As expected, we observed that both *PhCry2* and *PhPer* showed circadian expression in constant condition after entrainment to both LD and tides in WT (Figure S4A and B). As surprisingly observed by Oliphant et al., *PhCry2* and *PhPer* mRNAs also oscillate with an opposite phase after exposure to our entrainment conditions. While the *PhPer* expression pattern was clearly circadian, *PhCry2* showed a statistically significant 12-h period rhythm according to JTK cycle analysis, although the secondary peak was separated from the main peak by about 8 h. This is consistent with Oliphant et al., who could also observe a secondary *PhCry2* mRNA peak when varying the phase relationship between the LD and vibration cycles ^25^. In *PhCry2* −/− animals, *PhCry2* mRNA levels remained constantly low (Figures 1L and M). *PhPer* expression was also significantly disrupted as mRNA levels were significantly higher than in WT, except at CT18. A weak, but significant decrease in *PhPer*2 mRNA level was observed at CT12 and 18, but the phase of this weak apparent rhythm was completely distinct from that of the robust rhythms observed in WT animals. A similar progressive decrease in mRNA level was observed in mCry1 −/− mCry2 −/− double-mutant mice, perhaps as a result of prior mPer1 and mPer2 photic induction ^37^. Taken together, our results show that PhCRY2 is indeed part of the molecular circadian clock and is thus necessary for circadian behavior.

### *PhCry2* is an essential element of the circatidal clock

Next, we determined the role of PhCRY2 in the circatidal clock. We entrained animals to both LD and tidal (10.3 h high tide: 2.1 h low tide) cycles and monitored their behavior under DD and constant high tide (HT) conditions. As expected, WT and heterozygous animals showed strong rhythms of activity following the expected tides (Figure 2A and S2B). With both genotypes, low activity was observed at subjective low tide and activity peaked during the subjective high tide. In striking contrast, the homozygous animals did not show any rhythms of activity. All *PhCry2* −/− animal scored as arrhythmic (Figure 2B). A slight gene dosage effect was observed in heterozygous animals compared to WT, as the period of their circatidal behavior was significantly longer (Figure 2C, D and E). These results show that PhCRY2 is necessary to generate circatidal behavior.

**Figure 2:**
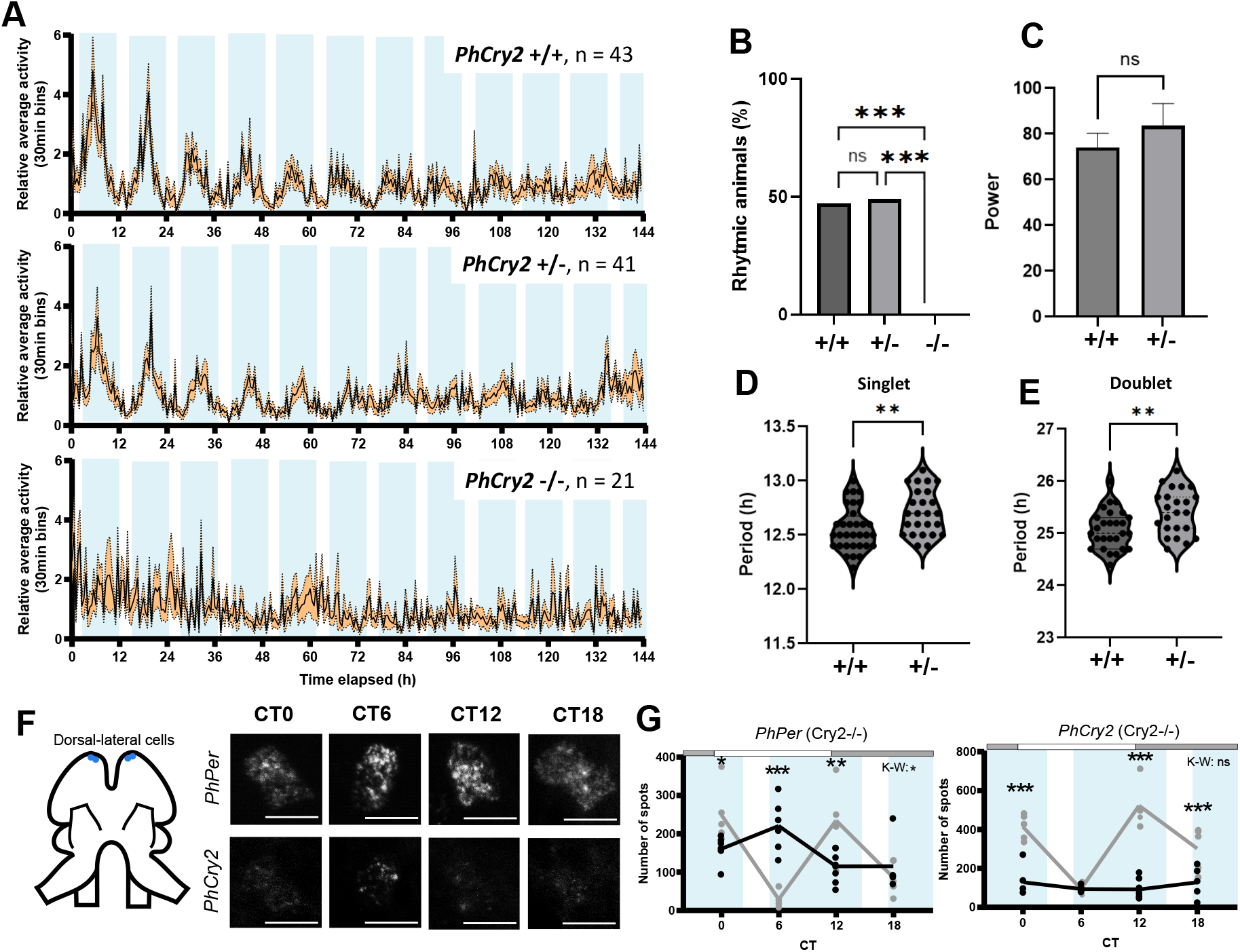
PhCRY2 is required to maintain circatidal rhythms in *P. hawaiensis*. A) Average relative swimming activity of wild-type (*PhCry2*+/+), heterozygous (*PhCry2*+/-) and homozygous (*PhCry2*−/−) animals recorded in DD and high tide after entrainment to a LD and tidal (10.3 h high tide: 2.1 h low tide) cycles. Blue shading represents the expected high tide. Standard error to the mean (SEM) is represented with the orange shading. B) Percentage of rhythmic animals under DD and high tide (Chi-square test, ***p <0.001; subsequent Chi-square test, +/+ vs +/-, p = 0.86, +/+ vs −/−, ***p <0.001 and +/- vs −/−, ***p <0.001). C) Rhythm power averages under DD and high tide (Welch’s t test, p = 0.416). D) Singlet period averages under DD and high tide (Welch’s t test, **p = 0.007). E) Doublet period averages under DD and high tide (Welch’s t test, **p = 0.005). F) HCR-FISH of *PhPer* and *PhCry2* mRNAs in circatidal dorsal-lateral clock neurons exposed to constant conditions after entrainment to 10.3:2.1 tides and 12:12 LD cycles in *PhCry2*−/− animals (*PhCry2* – mutation 1), as represented in Figure 1. G) As represented in Figure 1, quantification of the number of HCR-FISH spots in *PhCry2* −/− animals. In the mutant, there was an effect of time on gene expression for *PhPer*, (Kruskal-Wallis, *p = 0.0159), but not for *PhCry2* (Kruskal-Wallis, p = 0.545) (See Table S4 for post-hoc test p-value). Statistical differences between genotypes are indicated with * signs (Tukey’s multiple comparison, See Table S5 for exact p-value). See also Figure S2 and Table S4, S5 and S6

To assess the implication of *PhCry2* on molecular circatidal rhythms, we returned to HCR-FISH. The expression of *PhCry2* and *PhPer* was this time measured in the dorsal-lateral (DL) cells, which we found to be neurons as well (Figure S3B). The DL neurons entrain to vibration, exhibit ca. 12-h mRNA rhythms and are thus candidate circatidal oscillators ^25^. In WT, both genes showed robust 12-h rhythm of expression with low expression around the expected low tide (Figure S4C and S4D). In *PhCry2* −/−, as observed in circadian neurons, the expression of *PhCry2* remained constantly low (Figure 2F and G). Also, similarly to circadian neurons, *PhPer* expression was high and decreased over time, but did not show any sign of residual circatidal expression. In summary, our behavioral and molecular results demonstrate that PhCRY2 is an essential circatidal protein and is thus shared between the circadian and the circatidal systems.

### Loss of *PhPer* disrupts circadian rhythms

PhPER contains two Per-ARNt-Sim (PAS) domains and a Per-ARNt-Carboxy-terminal (PAC) domain. Three knock-out mutant lines were generated using CRISPR-Cas9 and a set of two gRNAs targeting an exon preceding those encoding the key functional domains (Figure 3A). The first mutant line (*PhPer* – mutation 1) harbored an 8 bp deletion combined with 2 added bp at the site targeted by the 5’ gRNA, and a 25-bp deletion with 4 added bp at the site of the 3’ gRNA. These indels thus caused a premature stop codon truncating PER from 1258 AA to 333 AA, and eliminating all key functional domains. The second mutant line (*PhPer* – mutation 2) had a premature stop codon following the deletion of 10 bp and the addition of 5 bp at the 5’ gRNA site, plus the deletion of 2 bp at the 3’ gRNA site, thus truncating PhPER to 317 AA. The last mutant line (*PhPer* – mutation 3), similarly to the others, had a premature stop codon and a truncated protein of 355 AA after a deletion of 9 bp with 3 extra bp at 5’ gRNA site, and the deletion of 7 other bp at the 3’ gRNA site. The absence of functional domains is predictive of a complete loss of function for all three mutations.

**Figure 3:**
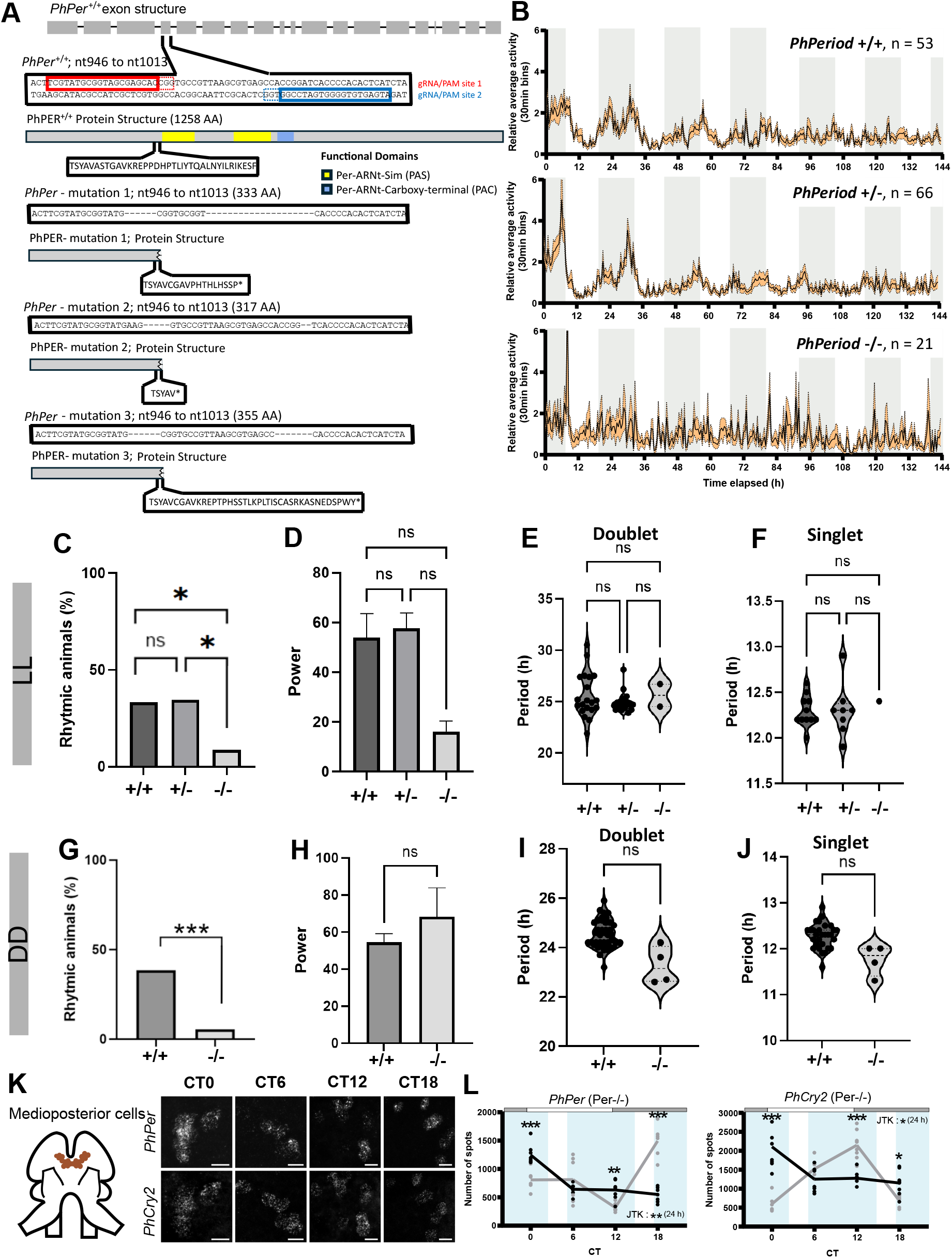
PhPER is required to maintain circadian rhythms in *P. hawaiensis*. A) Schematics of *PhPer* mutagenesis. Blue and red boxes indicate the gene sequences targeted by the gRNAs used for CRISPR/Cas9 mutagenesis. Three knock-out mutant lines with premature stop codons were obtained. The resulting mutant PhPER proteins are very similar, with slightly different C-terminal tails (see boxed amino acid sequences). All lines are missing key functional domains: the two Per-ARNt-Sim (PAS) (yellow) and the Per-ARNt-Carboxy-terminal (PAC) domains (blue). Thus, behavior data from the three strains were combined on this figure and on Figure 4. B) Average relative swimming activity of wild-type (*PhPer*+/+), heterozygous (*PhPer*+/-) and homozygous (*PhPer*−/−) animals recorded in constant light (LL) after entrainment to a LD cycle. Gray shading represents the subjective night. Standard error to the mean (SEM) is represented with the orange shading. C) Percentage of rhythmic animals in LL (Chi-square test, *p = 0.049; subsequent Chi-square test, +/+ vs +/-, p = 0.93, +/+ vs −/−, *p = 0.023 and +/- vs −/−, *p = 0.016). D) Rhythm power averages under LL (Kruskal-Wallis test, p = 0.137). E) Doublet period averages under LL (Kruskal-Wallis test, p = 0.544). F) Singlet period averages under LL (Kruskal-Wallis test, p = 0.746). G) Percentage of rhythmic animals under DD after LD entrainment (Chi-square test, ***p < 0.001; subsequent pairwise Chi-square test, ***p < 0.001). H) Rhythm power averages under DD (Mann-Whitney Smirnov test, p = 0.492). I) Doublet period averages under DD (Welch’s test, *p = 0.043). J) Singlet period averages under DD (Welch’s test, *p = 0.045). K) HCR-FISH to *PhPer* and *PhCry2* mRNAs in circadian medioposterior clock neurons exposed to constant conditions after entrainment to 10.3:2.1 tidal and 12:12 LD cycles in *PhPer* −/− animals (*PhPer* – mutation 1 and 3) as represented in Figure 1. L) As represented in Figure 1, quantification of the number of HCR-FISH spots in *PhPer* −/− animals. After testing the effect of time on the gene expression (*PhPer*, Kruskal-Wallis, ***p < 0.001, *PhCry2*, Kruskal-Wallis, ***p < 0.001), rhythm was tested using the JTK-cycle algorithm for 24-h periodicity (See Table S2 for exact p-value). Statistical differences between genotypes are indicated with * signs (Tukey’s multiple comparison, See Table S3 for exact p-value). See also Figures S5, S6 and Table S2, S3 and S6

Again, we first measured circadian behavior in LL. Both WT (*PhPer* +/+) and heterozygous animals (*PhPer* +/-) showed rhythmic behavior at the population level with higher activity during the subjective dark phase (Figure 3B and S5A). The homozygous mutant (*PhPer* −/−) population showed no clear rhythmic activity pattern. Accordingly, the percentage of rhythmic mutant animals was significantly lower than in *PhPer* +/+ and *PhPer* +/- populations (Figure 3C). Rhythm power of mutant animals was very low, but significance was not reached compared to wild-type and heterozygous, with only two animals showing rhythmicity (Figure 3D). Out of the two rhythmic *PhPer* −/− animals, only one had a period within the expected range (Figures 3E and F). When recorded in DD, homozygous animals were significantly less rhythmic than the WT with only four rhythmic animals out of 72 (Figures 3G to J and S6). Altogether, these results show that *PhPer* is critical to sustain circadian behavioral rhythms.

We then determined the effect of *PhPer* knock-out on the molecular circadian clock, with HCR-FISH staining on mutant animals entrained to LD and tidal cycles. As expected, both *PhCry2* and *PhPer* rhythms were disrupted in the circadian MP neurons, significantly decreasing over time instead of showing peaks of expression in the evening or night for *PhCry2* and *PhPer*, respectively (Figures 3K and L), reminiscent of the decay observed for *PhPer* mRNA in *PhCry2* −/− animals. As mentioned above, this could be because mRNA expression responds directly to light, and thus decreases progressively in DD. Altogether, these data show that loss of PhPER disrupts both molecular and behavioral circadian rhythms, as expected.

### *PhPer* sustains circatidal behavior as part of the circatidal clock

We then determined the role of *PhPer* in the circatidal clock. After entrainment to both LD cycle and tides (last high tide [LHT] 13:20), both WT and heterozygous animals showed lower average activity around the subjective low tides and higher activity around subjective high tides (Figure 4A and S5B). The average activity of animals lacking *PhPer* also showed this rhythmic pattern for 3 cycles, but it was much noisier and rhythm amplitude decreased rapidly. To determine if this residual rhythmicity was circatidal in nature, animals were entrained to a 6h-shifted tidal cycle (LHT 19h20) (Figure 4B and S5C). Again, *PhPer* mutants showed lower activity around the subjective low tides for 2, perhaps 3 cycles. Importantly, the phase of these residual rhythms was shifted by 6 hours, as in WT animals, confirming that they are entrained by tides (Figure 4C). However, the proportion of animals scoring as rhythmic was much lower in *PhPer* −/− mutants compared to WT and heterozygous animals (Figure 4D). For the few rhythmic mutant animals, power and period did not significantly differ from WT and heterozygotes (Figures 4E, F and G). These results show that PhPER sustains circatidal behavioral rhythms, although it might not be as critical as PhCRY2.

**Figure 4:**
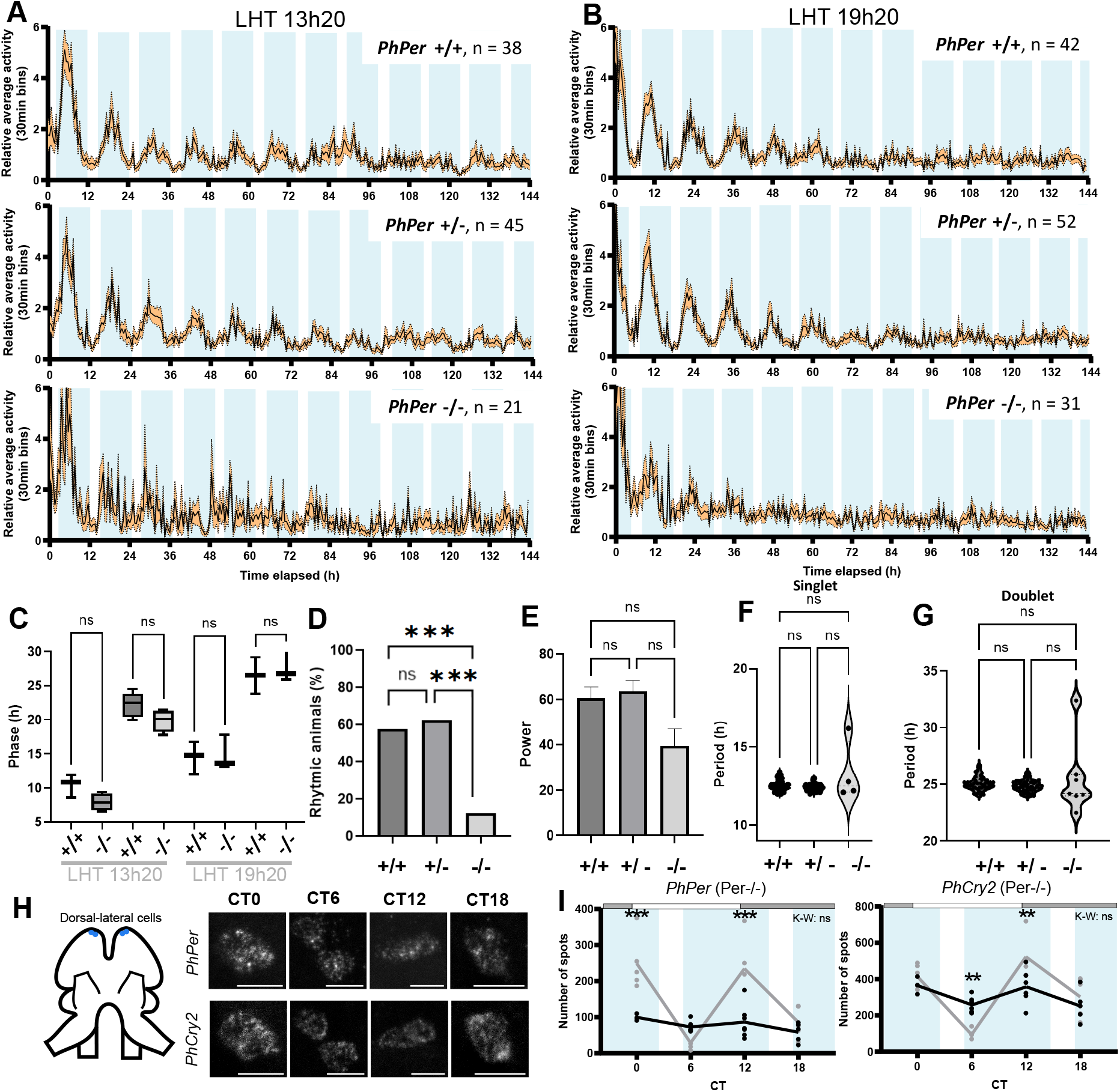
PhPER is required to maintain circatidal rhythms in *P. hawaiensis*. A) Average relative swimming activity of wild-type (*PhPer*+/+), heterozygous (*PhPer*2+/-) and homozygous (*PhPer*−/−) recorded in constant dark (DD) after entrainment to 10.3:2.1 tidal and 12:12 LD cycles as represented in Figure 2. The last high tide occurred at 13:20. B) Average relative swimming activity as represented in A. The tidal cycle was shifted by 6 hours; therefore the last high tide occurred at 19:20. C) Phase of maximum activity in *PhPer*+/+ and *PhPer*−/− under the two tidal regimes tested (One-way ANOVA, ***p <0.001 subsequent Sidak’s multiple comparison, LHT13:20 peak1, p = 0.380, LHT13:20 peak2, p = 0.347, LHT19:20 peak1, p = 0.999 and LHT19:20 peak2, p = 0.945). D) Percentage of rhythmic animals under DD and high tide. Tidal regimes were combined (Chi-square test, ***p <0.001; subsequent Chi-square test, +/+ vs +/-, p = 0.50, +/+ vs −/−, ***p <0.001 and +/- vs −/−, ***p <0.001). E) Rhythm power averages under DD and high tide (Kruskal-Wallis, p = 0.173) F) Singlet period averages under DD and high tide (Kruskal-Wallis, p = 0.300) G) Doublet period averages under DD and high tide (Kruskal-Wallis, p = 0.0602) H) HCR-FISH of *PhPer* and *PhCry2* mRNAs in circatidal dorsal-lateral clock neurons exposed to constant conditions after entrainment to 10.3:2.1 tides and 12:12 LD cycles in *PhPer* −/− animals (*PhPer* – mutation 1 and 3) as represented in Figure1. I) As represented in Figure 1, quantification of HCR-FISH spots per hemisphere across CT in *PhPer*−/− animals. (*PhPer*, Kruskal-Wallis, p = 0.174, *PhCry2*, Kruskal-Wallis, p =0.056). Statistical differences between genotypes are indicated with * signs (Tukey’s multiple comparison, See Table S5 for exact p-value). See also Figure S5 and Table S4, S5 and S6

We then determined the impact of the loss of PhPER on molecular circatidal rhythms in circatidal DL neurons. Neither *PhPer* nor *PhCry2* mRNAs showed statistically significant time-dependent variations in abundance (Figures 4H and I). Circatidal molecular rhythms are therefore clearly disrupted. However, we noticed slightly higher levels of both circadian mRNAs at the time at which expression is high in WT animals. Weak residual molecular rhythms might thus be present in PhPER−/− animals, which could explain the weak circatidal behavior rhythms that were observed. Overall, however, our data show that *PhPer* is part of the circatidal clock mechanism, although *PhPer* mutant animals can maintain weak circatidal behavioral rhythms in its absence, contrary to *PhCry2* −/− animals.

### *PhClk* is also required for circadian rhythms

CLK proteins are composed of a Beta-Helix-Loop-Helix (BHLH) domain required to bind to the E-Box and two PAS domains needed to bind to BMAL1 (Figure 5A). A previous study failed to identify CLK’s PAS-A coding sequences in the P. hawainesis genome or transcriptome. However, we did find two putative exons containing PAS-A coding sequences (Figure S7), indicating that PhCLK contains all three canonical domains. However, in the absence of transcriptomics support, we still cannot predict the full PhCLK sequence. To knock-out *PhClk* using CRISPR-Cas9, two gRNA were designed targeting the bHLH domain. Two mutant lines lacking all functional domains due to an early stop-codon in the BHLH domain were generated. The first line (*PhClk* – mutation 1) had a deletion of 27 bp with 4 extra bp inserted, generating a frameshift. The predicted truncated protein was 61 AA long instead of 2065 AA. The second line (*PhClk* – mutation 2) had a deletion of 80 nucleotides and generated truncated 42 AA protein.

**Figure 5:**
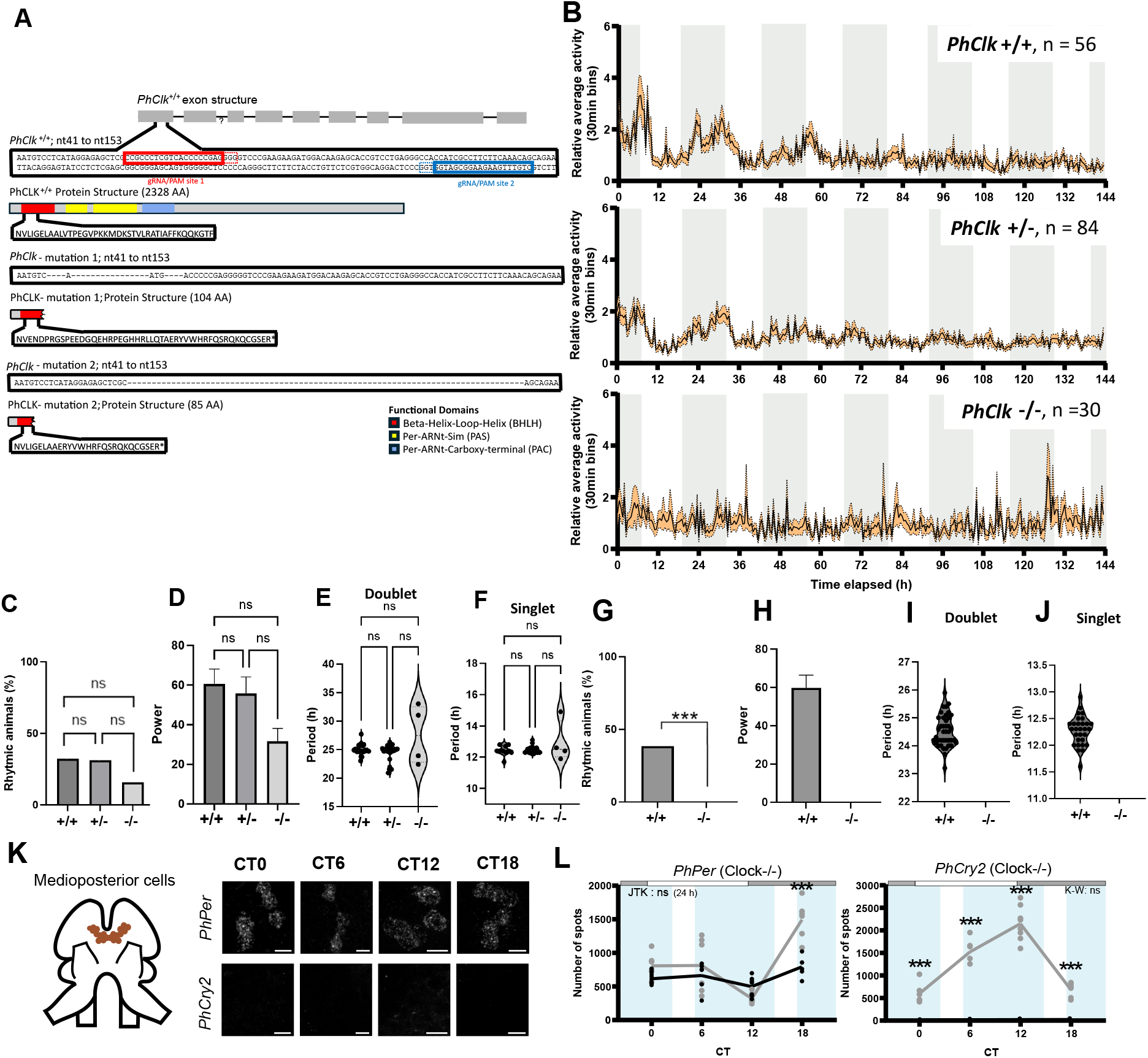
PhCLK is required to maintain circadian rhythms in *P. hawaiensis*. A) Schematics of *PhClk* mutagenesis. Blue and red boxes indicate the gene sequences targeted by the gRNAs used for CRISPR/Cas9 mutagenesis. Two knock-out mutant lines with premature stop codons were obtained. The resulting mutant PhCRY2 proteins are very similar, with slightly different C-terminal tails (see boxed amino acid sequences). Two mutant alleles were isolated, both missing part of the Beta-Helix-Loop-Helix (BHLH) domain (red), the Per-ARNt-Sim (PAS) domain (yellow) and the Per-ARNt-Carboxy-terminal (PAC) (blue). B) Average relative swimming activity of wild-type (*PhClk*+/+), heterozygous (*PhClk*+/-) and homozygous (*PhClk*−/−) animals recorded in constant light (LL) after entrainment to a LD cycle. SEM and expected LD and tidal cycles are represented as in Figure 1. C) Percentage of rhythmic animals in LL (Chi-square test, p = 0.157). D) Rhythm power averages under LL (Kruskal-Wallis test, p = 0.208). E) Doublet period averages under LL (Kruskal-Wallis test, p = 0.760). F) Singlet period averages under LL (Kruskal-Wallis test, p = 0.905). G) Percentage of rhythmic animals under DD after LD entrainment (Chi-square test, ***p < 0.001; subsequent pairwise Chi-square test, ***p < 0.001). H) Rhythm power averages under DD. I) Doublet period averages under DD. J) Singlet period averages under DD. K) HCR-FISH to *PhPer* and *PhCry2* mRNAs in circadian medioposterior clock neurons exposed to constant conditions after entrainment to 10.3:2.1 tidal and 12:12 LD cycles in *PhClk* −/− animals (*PhClk* – mutation 1) as represented in Figure1. L) As represented in Figure 1, quantification of the number of HCR-FISH spots in *PhClk* −/− animals. After testing the effect of time on the gene expression (*PhPer*, One-way ANOVA, *p = 0.013, *PhCry2*, Kruskal-Wallis, p = 0.528), rhythm was tested using the JTK-cycle algorithm for 24-h periodicity (See Table S2 for exact p-value). Statistical differences between genotypes are indicated with * signs (Tukey’s multiple comparison, See Table S3 for exact p-value). See also Figures S7, S8, S9 Table S2, S3 and S6

Both WT and heterozygous animals showed the expected circadian behavior pattern after LD-only entrainment, with higher activity during the expected dark phase (Figure 5B and S8A). By contrast, *PhClk* −/− animals did not show a rhythmic behavior pattern at the population level. Also, we observed fewer rhythmic animals in *PhClk* −/− compared to WT and heterozygous animals, although the difference did not reach significance (Figure 5C). Similarly, rhythm power was lower in *PhClk* −/− animals, though not significantly so (Figure 5D). Importantly, periods in the few weakly rhythmic animals were broadly distributed rather than clustered around 25.5 h, as previously observed in weakly rhythmic *PhBmal1* mutants ^24^ (Figures 5E and F). Circadian behavior thus appears to be disrupted. Moreover, *PhClk* was clearly required for circadian behavior under DD, as none of the knock-out animals were rhythmic under these conditions (Figures 5G to J and S9). Furthermore, HCR-FISH revealed that the molecular circadian clock is severely disrupted in the MP neurons of *PhClk* mutants. Neither *PhPer* nor *PhCry2* mRNA levels oscillated (Figures, 5K and L), with extremely low expression level of *PhCry2*. In summary, *PhClk* is required for the functioning of the circadian clock in *P. hawaiensis* and ensures proper circadian swimming activity.

### *PhClk* sustains circatidal behavior as part of the circatidal clock

We also probed the impact of loss of PhCLK on circatidal rhythms at the behavioral and molecular levels. In constant conditions following entrainment to both LD cycles and tides, both *PhClk* +/+ and *PhClk* +/- showed robust rhythmic behavior (Figures 6A and B and S8B). Animals lacking *PhClk* displayed a weakly rhythmic pattern for about 2 circatidal cycles that appeared entrained to tides, reminiscent of our observations with *PhPer* mutants. The proportion of rhythmic animals and the power of the rhythm were significantly lower in *PhClk* −/− compared to the other genotypes (Figures 6C and D). No significant difference was observed between genotypes for the singlet period (Figure 6E), although the doublet period of *PhClk* −/− was significantly different than WT and heterozygous animals (Figure 6F). Circadian phase was only quantified for the LHT 13h20 as only one *PhClk* −/− individual was rhythmic with the shifted tidal cycle (Figure 6G). No significant difference of phase was observed between mutants and WT, suggesting tidal entrainment of the weak residual rhythms. These results indicate that *PhClk* is necessary to sustain circatidal swimming activity.

**Figure 6:**
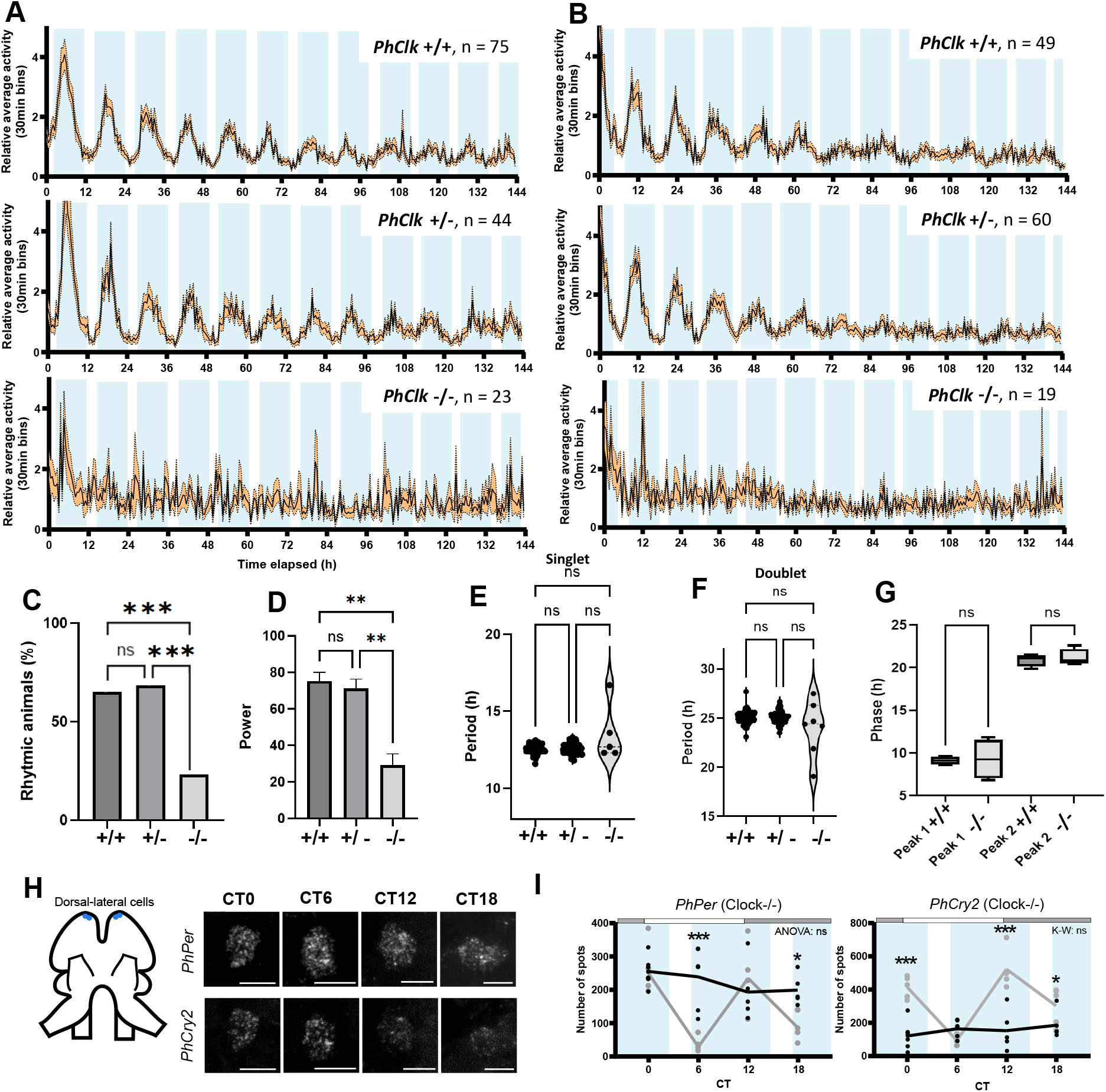
PhCLK is required to maintain circatidal rhythms in *P. hawaiensis*. A) Average relative swimming activity of wild-type (*PhClk*+/+), heterozygous (*PhClk*+/-) and homozygous (*PhClk*−/−) animals recorded in constant dark (DD) and high tide after entrainment to a LD cycle and tidal (10.3 h high tide: 2.1 h low tide) as represented in Figure 2. The last high tide occurred at 13:20. B) Relative average swimming as represented in A. The tidal cycle was shifted by 6 hours, so the last high tide occurred at 19:20. C) Percentage of rhythmic animals under DD and high tide. Tidal regimes were combined (Chi-square test, ***p <0.001; subsequent Chi-square test, +/+ vs +/-, p = 0.564, +/+ vs −/−, ***p <0.001 and +/- vs −/−, ***p <0.001). D) Rhythm power averages under DD and high tide (Kruskal-Wallis, **p = 0.002; subsequent Dunn’s multiple comparison, +/+ vs +/-, p = 1, +/+ vs −/−, **p = 0.001 and +/- vs −/−, **p = 0.005). E) Singlet period averages under DD and high tide (Kruskal-Wallis, p = 0.673). F) Doublet period averages under DD and high tide (Kruskal-Wallis, p = 0.350). G) Phase of maximum activity in *PhClk*+/+ and *PhClk*−/− in the two tidal regimes tested (One-way ANOVA, ***p <0.001 subsequent Sidak’s multiple comparison, LHT13:20 peak1, p = 0.982, LHT13:20 peak2, p = 0.946). H) HCR-FISH to *PhPer* and *PhCry2* mRNAs in circatidal dorsal-lateral neurons exposed to constant conditions after entrainment to 10.3:2.1 tidal and 12:12 LD cycles in *PhClk* −/− animals (*PhClk* – mutation 1) as represented in Figure1. I) As represented in Figure 1, quantification of HCR-FISH spots in *PhClk*−/− animals (*PhPer*, One-way ANOVA, p =0.370, *PhCry2*, Kruskal-Wallis, p =0.380). Statistical differences between genotypes are indicated with * signs (Tukey’s multiple comparison, See Table S5 for exact p-value). See also Figure S8 and Table S4, S5 and S6

At the molecular level, both *PhPer* and *PhCry2* were detected in circatidal DL neurons of *PhClk* mutants (Figure 6H and I) but showed no time-dependent changes in expression. *PhPer* was at a constant high level compared to WT, while *PhCry2* was constantly low. This shows that *PhClk* is also required for the functioning of the circatidal molecular clock.

### *PhBmal1* also controls circadian and circatidal expression, but PhBMAl1 and PhCLK functions differ in circatidal neurons

Our previous work showed that *PhBmal1* is required to maintain both circadian and circatidal activity in *P. hawaiensis*, but its impact on the molecular circadian and circatidal clock was not tested ^24^. We thus performed FISH-HCR on *PhBmal1* knock-out animals. In the circadian MP neurons, the circadian expression of *PhPer* and *PhCry2* was deeply disrupted (Figures 7A and B). The expression of *PhPer* decreased over time whereas *PhCry2* expression was low at all time points. Nevertheless, *PhCry2* showed a weak rhythm with a completely abnormal phase. We therefore conclude that PhBMAL1 is necessary for proper circadian molecular rhythms. In the circatidal DL neurons, neither repressor genes showed circatidal expression (Figures 7C and D). This confirms that BMAL1 is part of the circadian and the circatidal clocks systems in *P. hawaiensis*.

**Figure 7.**
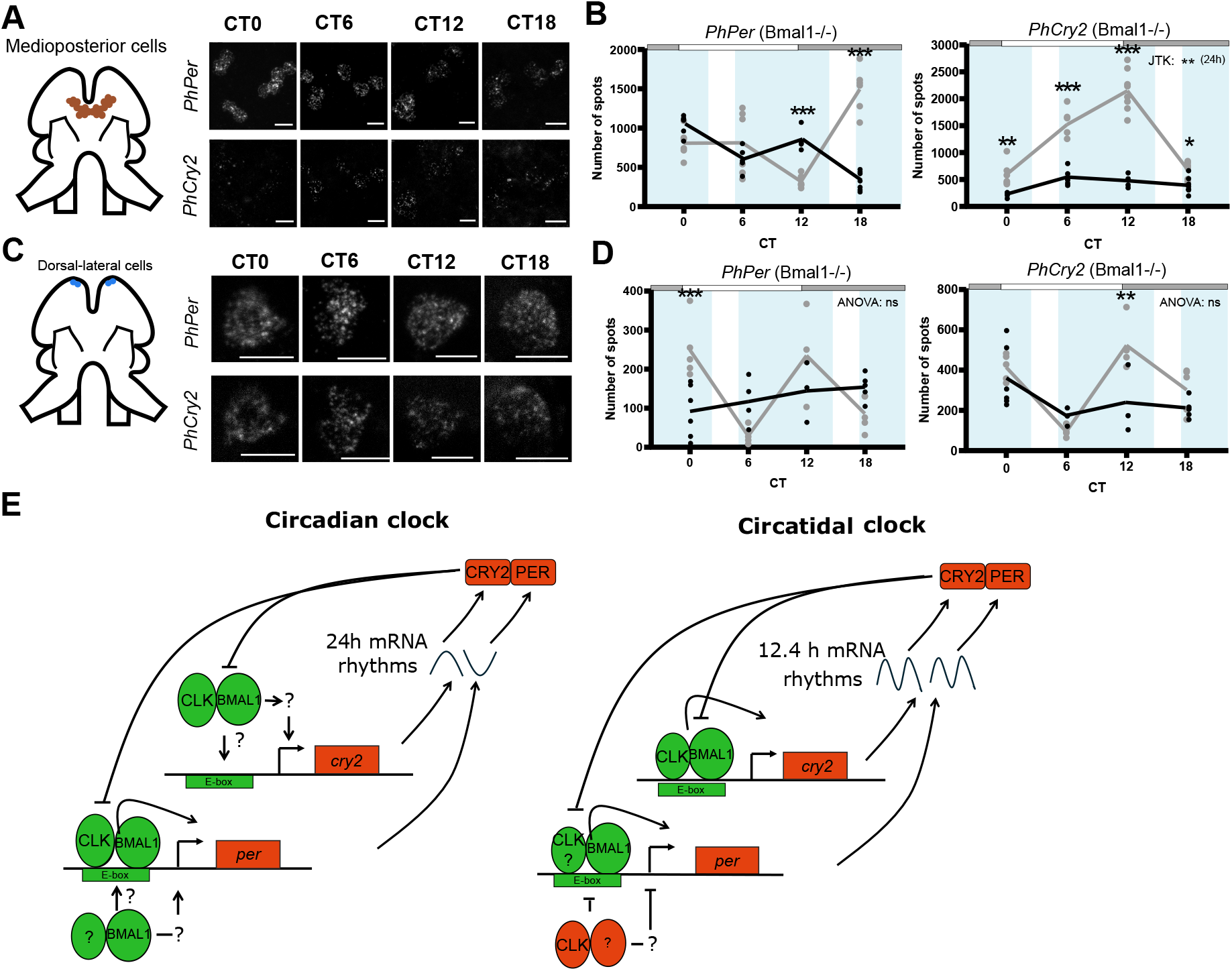
PhBMAL1 is part of the molecular circadian and circatidal clocks in *Parhyale*. A) HCR-FISH to *PhPer* and *PhCry2* mRNAs in circadian medioposterior clock neurons exposed to constant conditions after entrainment to 10.3:2.1 tidal and 12:12 LD cycles in *PhBmal1* −/− animals as represented in Figure 1. B) As represented in Figure 1, quantification of the number of HCR-FISH spots in *PhBmal1* −/− animals. After testing the effect of time on the gene expression (*PhPer*, One-way ANOVA, ***p < 0.001, *PhCry2*, One-way ANOVA, **p = 0.002), rhythm was tested using the JTK-cycle algorithm for 24 h (See Table S2 for exact p-value). Statistical differences between genotypes are indicated with * signs (Tukey’s multiple comparison, See Table S3 for exact p-value). C) HCR-FISH of *PhPer* and *PhCry2* mRNAs in circatidal dorsal-lateral clock neurons exposed to constant conditions after entrainment to 10.3:2.1 tides and 12:12 LD cycles in *PhBmal1* −/− animals as represented in Figure 1. D) As represented in Figure 1, quantification of HCR-FISH spots in *PhBmal1*−/− animals. (*PhPer*, One-way ANOVA, p =0.376, *PhCry2*, One-way ANOVA, p =0.098). Statistical differences between genotypes are indicated with * signs (Tukey’s multiple comparison, See Table S5 for exact p-value). E) Circadian and circatidal clock models for *P. hawaiensis*. Activators are in green, and repressors in red. Based on our mutant analysis, we propose that the two clocks are wired differently, transcriptionally. A key difference is PhCLK’s role in *PhPer* regulation. In the circadian clock, PhCLK promotes *PhPer* transcription. However, in the circatidal clock, it is predominantly a *PhPer* repressor. It might do so through direct binding to the *PhPer* promoter, or by promoting the expression of an unknown transcriptional repressor. *PhBmal1* promotes *PhPer* expression in circatidal neurons, either with the help of *PhClk*, or a different partner. In the circadian neurons, *PhPer* and *PhCry2* mRNAs cycle in antiphase. This could be because *PhCry2* is indirectly regulated by *PhBmal1* and *PhClk*, through posttranscriptional regulation, or perhaps because PhBMAL1 also directly or indirectly regulates *PhPer* with an unknown partner in addition to *PhClk*. See main text for more discussion. See also Table S2, S3 S4, S5

As expected, since BMAL1 is a transcriptional activator, *PhPer* and *PhCry2* mRNAs remained constantly at a low range in both circadian and circatidal neurons, never reaching WT peak expression. This was also the case in brain circadian neurons of *PhClk* mutants (Figure 5K and L), as expected since CLK and BMAl1 work as dimers. In stark contrast, while *PhCry2* mRNA was as expected also low, *PhPer* mRNA was constantly high in circatidal neurons of *PhClk* mutant animals (Figure 6H and I). PhCLK thus represses, rather than activate, *PhPer* expression specifically in circatidal neurons. This reveals a surprising difference in the transcriptional wiring of core clock genes between circadian and circatidal neurons, and a novel PhBMAL1-independent function for PhCLK (Figure 7E).

## Discussion

We previously showed that *PhBmal1* is necessary for circatidal behavior in *P. hawaiensis*, in addition to its canonical role in the circadian clock ^24^. With the present study, we find that there is considerable overlap between circadian and circatidal clock mechanisms. Using CRISPR-Cas9 genome editing, we demonstrate that the three other core circadian genes are required to sustain circatidal behavior. This is in stark contrast to previous studies in *E. pulchra* and *A. asahinai* ^19–22^. It is possible that the absence of RNAi phenotypes when targeting Per, Clk and Cry2 is the result of incomplete suppression of gene expression. In support of this idea, RNAi to EpBmal1 decreased the percentage of rhythmic animals, but this phenotype was much weaker than the complete disruption observed in *PhBmal1* knock-out animals ^24^. Another possibility is that circatidal timekeeping mechanisms differ across marine organisms, resulting from convergent evolution or divergent evolution of a common proto-circatidal clock. The only previous study made at the cellular level gives some support to these possibilities: in putative circatidal cells of *E. pulchra*, only EpTim showed circatidal expression, whereas EpPer and EpCry2 were either arrhythmic or displaying a circadian expression pattern ^25^. In contrast, in *P. hawaiensis*, which lacks Tim, both *PhPer* and *PhCry2* showed circatidal expression. Also, a recent study in which a CRISPR-Cas9 Per knockout was generated in *A. asahinai* concluded that PER plays no role in circatidal rhythms in this insect. However, this conclusion was based on a single rhythmic animal out of 10 monitored, and no molecular rhythms were measured ^38^.

In *P. hawaiensis*, we found that loss of both PhCRY2 and PhBMAL1 completely disrupted molecular and behavioral circatidal rhythms ^24^. On the other hand, in both *PhPer* −/− and *PhClk* −/− animals, residual circatidal behavioral rhythms are observed, even though molecular rhythms are severely disrupted. This demonstrates that PhPER and PhCLK are important cogs of the circatidal clocks, although we surmise that residual molecular rhythmicity is present in their absence, at least in the few animals that exhibited circatidal behavior. It is possible that another bHLH/PAS transcription factor can weakly substitute for PhCLK. PhHIFα, in particular, comes to mind, since in mammals it can dimerize with BMAL1 and has been implicated in the entrainment of circadian clock by O_2_ levels ^39–42^. Weak rhythms might be possible in the absence of PhPER, because repressive cryptochromes are the dominant circadian repressors in most species, while PER have supporting functions, such as promoting CRY nuclear entry ^29,43^. This was confirmed by our luciferase assay, where PhCRY2 alone is able to repress PhBMAL1:mCLK transactivation, while PER was unable to do so.

A recent study revealed that while *PhPer* and *PhCry2* mRNAs cycle in phase inside circatidal neurons, their oscillations are unexpectedly antiphasic in circadian neurons ^25^. We confirmed these observations under our tidal entrainment protocol. This suggests that the transcriptional network linking different clock genes is wired differently in circadian and circatidal neurons. We therefore examined closely the impact of core clock gene mutations on *PhPer* and *PhCry2* mRNA levels in both circadian and circatidal neurons. In both neuron types, the loss of *PhPer* led to moderately high mRNA levels of *PhCry2*, and the loss of *PhCry2* to moderately high *PhPer* mRNA levels. This is consistent with a repressive role for both CRY2 and PER. Note that the loss of core circadian repressors does not result in peak mRNA levels in *Drosophila*, because interlocked feedback loops are also disrupted by the loss of core clock repressors, thus reducing DmCLK levels ^44,45^. Low *PhPer* mRNA levels were observed in *PhPer* −/− mutants, and low *PhCry2* mRNA levels in *PhCry2* knock-outs. Since our mutations create premature stop codons, these low mRNA levels are probably the result of nonsense-mediated mRNA decay ^46^.

Our FISH-HCR and cell culture assays thus support a rather standard repressive role for PhPER and PhCRY2 in both circadian and circatidal clocks. They also support the expected activator role of *PhBmal1*, since *PhCry2* and *PhPer* mRNA levels are low in both clock neuron types of *PhBmal1* mutants. In circadian neurons, the loss of *PhClk* also led to low mRNA levels, supporting its canonical role as a circadian transcriptional activator along with BMAL1. However, *PhCry2* levels were particularly low in MP neurons, essentially undetectable. This was not linked to a technical issue, since we detected *PhPer* mRNAs in these circadian neurons and *PhCry2* mRNAs in the circatidal DL neurons of the same brains. This suggests that PhBMAL1 might be able work with a different partner than PhCLK to weakly activate and therefore maintain constant low transcript levels of *PhPer*, but is unable to do so for *PhCry2* in circadian MP neurons. As mentioned above, PhHIF1α could substitute for PhCLK. Also, HIF1α has been hypothesized to bind to the non-canonical EBox2 of mPeriod2 ^47,48^. Therefore, we propose that *PhPer* and *PhCry2* genes are activated by different sets of transcription factors in circadian neurons (Figure 7E). This could contribute to the phase difference in expression of the two repressors. Other possibilities include differences in mRNA stability, and PhCLK/PhBMAl1 controlling an intermediate positive *PhCry2* regulator (Figure 7E), which would work similarly to RORα or PDP1ε in promoting antiphasic expression of mammalian Bmal1 and D*mClk*, respectively ^6,8,9,49,50^.

In circatidal neurons missing *PhClk, PhPer* levels were constantly at peak levels while *PhCry2* levels were constantly low. This is rather unexpected. It indicates that *PhClk* represses *PhPer* expression independently of *PhBmal1* (Figure 7E). This is a striking deviation from its regular function in the circadian clock mechanism. It will be interesting to determine whether PhCLk represses *PhPer* as a homodimer, or with a novel partner, and whether its impact on *PhPer* expression in circatidal neurons is direct or mediated by an intermediate repressor. Critically, PhCLK represses *PhPer* expression specifically in circatidal neurons.

In conclusion, transcriptional control within the core circadian and circatidal clocks is wired differently (Figure 7E). However, based on the surprisingly antiphasic oscillations of *PhPer* and *PhCry2* mRNAs in circadian neurons, one would have expected the major deviation from the canonical function of core clock genes to occur in these neurons, rather than in circatidal neurons. Obviously, deeper studies will be necessary to resolve this apparent paradox. However, the identification of a novel function for *PhClk*, specific to circatidal neurons, could be critical to understanding how circadian and circatidal clocks oscillate with such different periods. Among the three historical models for circatidal rhythms, our observations, combined with those of Oliphant et al. ^25^, support the model proposed by Naylor ^14^, of separate circatidal and circadian clocks, even though they use the same core clock genes. However, because of their overlapping mechanisms, we cannot exclude entirely the Enright model of a plastic clock ^15^. It is possible that during development, clock neurons adopt different transcriptional wiring based on the sensory neurons they are coupled with, and thus the environmental input they receive: tidal or diurnal. We note that the period of core clock gene expression changes as a function of the presence of tides in oysters, but the mechanism underlying this plasticity is not known ^51^. We also note that the same species of freshwater snail can adopt more circadian or circatidal phenotypes (behaviorally and molecularly), depending on whether the animals live within or outside of the tidal zone of the Kiso River in Japan^52,53^. In *P. hawaiensis*, it will be very interesting to determine what dictates the periodic fate of clock neurons, and whether this fate could depend on environmental factors.

Regardless, our results reveal that in *P. hawaiensis*, the evolutionary solution to the conundrum of simultaneously tracking tidal and diurnal environmental oscillations that are almost, but not exactly, harmonics, is to use the same set of core clock genes in two spatially distinct neuronal clocks, each having different autoregulatory feedback loops to set their specific periodicity. This spatial separation of the circadian and circatidal clocks likely underlies *P. hawaiensis* ability to integrate diurnal and tidal cues, rendering it capable of adapting its behavior and physiology to the different phase relationships of the two environmental cycles. It also likely permits plastic behavioral responses to different tidal regimens. Indeed, *P. hawaiensis* lives in a broad range of locations that experience different tidal patterns ^54^ : semi-diurnal (the most common 12.4h tidal cycle), mixed (two tides per 24.8h, of unequal duration and amplitude) and diurnal (24.8-h tidal cycle). This work improves our understanding of the complex mechanisms by which an organism copes with complex environmental periodicities. The extent to which this mechanism applies across marine organisms remains to be determined. Given the conservation of the circadian clock, these rules could prove universal.

## Material and Method

### Animal husbandry

The Chicago F-strain *Parhyale hawaiensis* strain was used for our study. The animals were maintained in Pyrex aquariums with lids containing 30 psu artificial sea water (ASW) and ∼1 cm of crushed coral substrate. The animals were reared at a temperature of 25°C under 12:12 h LD cycles with ZT0 occurring at 08:00 in the morning. Wild type (WT) and *PhBmal1* mutants ^24^ were fed baby carrots. *PhCry2, PhPer* and *PhClk* mutants were fed with a mix of ground TetraMin PLUS Tropical Flakes (Tetra, Melle, Germany), ground OSI Spirulina Flake Fish Food (Ocean Star International, Snowville, UT), ground Hikari Wheat Germ Floating Pellets for Pets (Hikari, Japan), ground Tubifex (Hagen Group, Quebec, Canada), and Kelp granules (Starwest Botanicals, Sacramento, CA). The dry food was mixed in 50 mL falcon tubes containing ASW supplemented with 50 µL of American Marine Selcon Vitamin Supplement (American Marine Inc., Ridgefield, CT) and 100 µl of KENT Marine Zoe Marine Vitamin (KENT Marine, Franklin, WI).

### CRISPR-Cas9-mediated mutagenesis

Mating pairs were isolated a day prior to embryo removal. Females were then lightly anesthetized using 0.02% clove oil in ASW. The collection of single-cell fertilized eggs was achieved as previously described ^24^. Fertilized eggs were stored in a tissue culture dish containing filtered ASW water prior to injection. Approximately 40-60 picoliters of injection mixture was injected into one-cell embryos or both cells of two-cell embryos, as previously described ^55^. The injection mixture contained 100 ng/µL or 162 ng/µL of each chemically protected guide RNA, 333 ng/µL of NLS-Cas9 protein (Synthego Co., Redwood City, CA), and 0.05% phenol red (Sigma-Aldrich). Both injection mixture concentrations generated knockouts. For each gene of interest, two guide RNAs were designed to target a functional domain or a region prior to a key functional domain. Injected embryos were then transferred to 60 mm culture dishes filled with filter-sterilized ASW and incubated at 28°C under a 12:12 h LD cycles. Embryos were transferred on a daily basis to a new culture dish containing ASW. After hatching, juvenile animals were reared individually in a custom multi-compartment insert placed inside a lidded plastic container filled with ASW. Each compartment had a mesh-covered opening that allowed water exchange with the shared bath while preventing mixing and aggression/cannibalism between individuals. Once injected animals reached sexual maturity, they were paired with other injected or wild-type individuals of the opposite sex to generate heterozygous offspring (F1) with fixed germline mutations. Heterozygotes F1 were then reared in a separate multi-compartment tank until sexual maturity. After the screening of mutations (see below), F1 males having targeted genotype were individually selected and transferred individually to a larger aquarium (20×10×10 cm) with at least 4 wild-type females. These tanks contained ∼2 cm of crushed coral substrate and were fed with the food mix described above.

For the screening of mutations, adult animals were anesthetized using clove oil in ASW, after which two to three pleopods were removed and digested in 250 µL of lysis buffer (100 mM Tris, 5 mM EDTA, 200 mM NaCl, 0.2% SDS) containing 100 µg/mL of proteinase K at 50°C for 48 h. The tubes were centrifuged at 13000 rpm at 4°C for 10 min to pellet the debris. DNA was precipitated by adding 250 µL of isopropanol to the supernatant followed by centrifugation for 3 minutes at room temperature. The pellet was then washed with 500 µL of 70% ethanol, dried on a heat block at 55°Cand resuspended in 20 µL of milliQ water. PCRs were performed using Standard Taq DNA polymerase (New England BioLabs) as indicated by the manufacturer. PCR primers are listed in Table S1. The amplification program consisted of 35 cycles of denaturation at 95°C for 30 s, 30s annealing (54°C for *PhCry2* primers, and 56°C for *PhPer PhClk* primers) and elongation at 68°C for 20 s. PCR products were run on a 3% agarose gel and the genotype was confirmed by gel purification of the products followed by sanger sequencing.

### Entrainment and behavioral monitoring under constant conditions

Tidal entrainment was performed as previously described ^24^ with cycles of 10.3 hr of high tide and 2.1 hr of low tide. Low tide was simulated when water was transferred to a reservoir using a draining pump. For high tides, water was pumped back into the aquarium containing the animals. Animals were entrained to tides for a minimum period of 30 days prior to the recording of behavior. After at least five days of undisturbed entrainment (i.e. no water changes or feeding), males and females were individually loaded into polystyrene vials containing ∼2 mm of crushed coral substrate and 30 psu ASW. The water height was adjusted to equal their original aquaria level, thereby maintaining a constant high tide. ParaFilm was used to close the vial and limit evaporation. *Drosophila* Activity Monitors (DAM) (LAM25, TriKinetics, Waltham MA) placed in a temperature and light controlled incubator (I36LL, Percival Scientific, Iowa) were used for behavioral recordings. Following a day of acclimation, swimming and roaming activity were recorded for a duration of five days with recordings conducted at one-minute intervals. After LD-only entrainment, behavior was recorded under either constant light (LL) or constant darkness conditions (DD). The behavior after tidal and LD entrainment was recorded in DD. A minimum of three sets of experiments were conducted for each condition and for each genotype.

For most behavioral studies, the mutant colonies were of mixed genotypes, containing wild type (WT), heterozygous and homozygous mutant animals. Their genotypes were determined after recording as described above for mutant screening. For DD experiments and FISH-HCR experiments (see below), mutant colonies containing almost exclusively homozygous animals were used. These animals were nonetheless genotyped after behavioral monitoring or prior to brain dissection.

### Behavior analysis and statistical analysis

The recorded one-minute beam-crossing was averaged over 30-minute bins using DAMFileScan114 software (TriKinetics, Waltham MA). The rhythm analysis was performed with the FaasX software (https://neuropsi.cnrs.fr/en/cnn-home/francois-rouyer/faas-software/, courtesy of F. Rouyer, Centre National de la Recherche Scientifique, Gif-sur-Yvette, France) ^56^. Autocorrelation and χ^2^ periodogram analysis (“power” ≥ 10 and “width” ≥ 1h) were used to determine whether an animal was rhythmic. The filter for high frequencies was on. As previously described, both “vertical swimming” and “roaming” behaviors were monitored ^24^. Animals were scored as rhythmic if vertical swimming and/or roaming were significantly rhythmic. For each animal, the period and rhythm power of the most robustly rhythmic of these two behaviors was used for subsequent analysis. For the population activity plots, behavior values were normalized for each animal by dividing each 30 minutes bin by its mean activity. Plots were generated using GraphPad Prism 10. Statistical comparisons of the power and the period were made using the same software. Outliers were identified and removed using a ROUT test with Q=1%. The normality was assessed using Shapiro-Wilk test. If the normality was respected, one-way ANOVA test was used followed by Tukey’s for multiple comparisons. Otherwise, the Kruskal-Wallis test was used, followed by Dunn’s multiple comparisons test. In instances where the comparison was between two groups, normal dataset was analyzed using Welch’s test for normal distribution. Otherwise, Mann-Whiney test was used. Statistical significance is demonstrated as ***p < 0.001; **p < 0.01; *p < 0.05.

### Brain sampling and processing

Animals were entrained to tides under LD cycles as described above. After LHT 12h10, five animals per sampling point were loaded separately in vials in the DAM system as described before (their behaviors are shown on figure S10). The animals were in DD after the light was switched off at 20h00. The following day, the sampling process was initiated at 8h00. First WT animals were sampled every two hours over 24 hours (13 sampling points). For the mutants, four sampling points spaced by six hours were chosen based on WT results. The following strains were used: *PhCry2* mutation 1, *PhPer* mutations 1 and 3 and mutation 1 for *PhClk*. For the brain sampling, the animals were sacrificed by immersion in 4°C ASW for 20 minutes on ice. Heads were removed and placed in 4% PFA prepared in 3X PBS for 4 hours with rotation at 4°C. The heads were then washed with PBS-Tween 0.1% for 5 minutes three times. Brains were dissected in PBST using sharpened forceps. The brains were then dehydrated using a methanol dilution series (50%, 70% and 90% methanol in PBST) for five minutes each wash, followed by two brief washes in 100% methanol. Then brains were stored for a minimum of 24 hours at −20°C before staining.

### HCR™ RNA-FISH

Prior to staining, the brains were rehydrated using serial dilutions of methanol (5 min in 70%, 50% and 25%) followed by 1X10min and 2X5min washes in PBST. They were then incubated overnight in 300 µL of detergent solution ^57^ (50mM Tris, 1mM EDTA, 150mM NaCl, 0.5% Tween, 1% SDS) at 4°C. The following day, the brains were pre-hybridized in 200µl hybridization buffer (Molecular Instruments Inc) at 37°C for 3 hours. The brains were then incubated in 100 µl of hybridization buffer containing 20nM *PhCry2, PhPer* and *PhClk* probes at 37°C. To determine neuronal identity, cells were co-stained using *PhPer* and PhElav probes (Figure S3). HCR™ probes V3.0 were designed and synthesized by Molecular Instruments Inc, USA. After 48 h of incubation, the brains were washed 1×5min, 1×10min and 3×15min with 500µL of wash buffer (Molecular Instruments Inc) at 37°C and then 3X5min in 5XSSC supplemented by 0.1% Tween-20 at room temperature (RT). Pre-amplification consisted of 2×15min washes in 500µL of amplification buffer (Molecular Instruments Inc) at RT. Fluorescent hairpins were snap-cooled and added to 100µl of amplification buffer at a concentration of 60 nM. The brains were incubated overnight in the hairpin preparation at RT. Washes consisted of 2X5min, 2X30min and 1X5min in 500µL of 5XSSCT. The brains were then incubated in 50% glycerol in PBS containing 2µg/ml of DAPI for 1 hour at RT followed by 20 minutes in 70% glycerol. Brains were mounted in VectaShield Antifade Media (Vector Laboratories, USA) between two coverslips spaced using a Grace Bio-Labs SecureSeal™ imaging spacer. Images were acquired with a LSM 900 Zeiss confocal microscope at 1µm z-stack intervals.

### Image analysis

Image analysis was conducted using ImageJ software. Prior to quantification, the images were smoothed and each cell group was isolated. The number of FISH dots, which reflect the expression of the gene, was counted using the RS-FISH ImageJ plugin ^58^. The normality of the dataset was assessed using a Shapiro-Wilk test. If a normal distribution was observed, a one-way ANOVA was used to determine if significant variations occurred though time, otherwise a Kruskal-Wallis test was used. When a statistically significant effect was observed, rhythm analyses were conducted in R (v.4.4.1) ^59^ using JTK CYCLE in the package MetaCycle ^60^ for 24 h periods. P-values were adjusted using the Bonferroni method. For 12-h rhythms, the statistical difference between peaks and trough were calculated using post-hoc tests.

### Transcriptional assay in HEK cells

The TK-E54 plasmid ^61^ harboring 3 E-Boxes driving the expression of the firefly luciferase reporter, the CMV β-galactosidase plasmid (Clontech: PT2004-5) ^28^ containing the LacZ gene under the control of the CMV promoter and the *mClk*^*62*^ cDNA under the CMV promoter in pcDNA3.1 were all provided by D. Weaver (University of Massachusetts Chan Medical School). The *PhBmal1* plasmid has been previously described ^24^. *PhPer* and *PhCry2* were produced by cutting *mClk* plasmid using Xhol and EcoRI. gBlocks (Twist Biosciences) were designed to contain a start codon, a Flag tag (*PhPer*) or V5 tag (*PhCry2*) and the whole *PhPer* or *PhCry2* sequences flanked by Xhol and EcoRI. Sequences used were based on Hunt et al. in silico reconstruction ^26^. *PhCry2*−/− plasmid was generated by integrating an early stop-codon into *PhCry2* plasmid to mimic the mutations generated in vivo. To do so, the *PhCry2* plasmid was amplified through PCR using primers designed to introduce premature stop codon. As part of this amplification, the downstream portion of the gene was removed, and a homologous overhang was included for later re-ligation of the plasmid. The resulting *PhCry2* −/− is missing a part of the photolyase domain and the FAD binding domain, as the KO lines used in this paper. Following the amplification, Dpn1 digestion (New England Biology) was achieved following the protocol outlined by manufacturers. Following digestion, a DNA cleanup step was performed using the QIAquick PCR purification kit (Quiagen). The cleaned *PhCry2*−/− pcDNA3.1 amplicon was then religated through homologous assembly using the NEB Gibson Assembly kit (New England Biology) generating the finalized construct.

HEK-293T cells were plated at 5.10^5^ cells per well in 6 well plates (Thermo Fisher Scientific) and were transfected 24-h later using Lipofectamin 2000 Transfection Reagent from Invitrogen (Waltham, MA). A total of 1600 ng of DNA was transfected per well (30 ng of TK-E54, 100 ng of *PhBmal1, mClk* and CMV β-gal and different increasing concentration of repressors (i.e. 0, 10, 50, 200 and 500 ng of *PhCry2* and/or *PhPer* or *PhCry2*−/−), supplemented with empty pcDNA3.1 as needed. After transfection, plates were incubated 48 h at 37°C and 5% CO_2_. They were then washed with ice-cold PBS and incubated 15 min in 350 µL of Reporter Lysis Buffer (Promega). Cell lysates were removed from the well and spun down at 13000 rpm. The supernatant was used immediately for assays or stored at −80°C.

Following the manufacturer protocol for the β-Galactosidase assay, 50 µL of the lysate was mixed with 50 µl Assay Buffer (Promega) in a 96 well plate (Corning) and incubated for 30 minutes at 37°C or until a faint yellow color appeared. The reaction was stopped using 150 µl of 1M Sodium Carbonate and the absorbance was read at 420 nm using a SpectraMax iD5 microplate reader (Molecular Devices, San Jose, CA). For the luciferase assay, 20 µl of the lysate was added to 20 µl of D-luciferin mix (0.132mg/mL D-luciferin (Sigma), 20mM Tricine, 2.67mM MgSO4, 0.1mM EDTA, 33.3mM DTT, 530uM ATP (Sigma), 270uM acetyl coenzyme A (Sigma), 265uM 4MgCO3Mg(OH)2) and the luminescence was measured using the same microplate reader. The β-Galactosidase signal was used to normalize the firefly luciferase signal.

## Acknowledgments

We thank Erica Kwiatkowski for generating the pAc-*PhPer* and pAc-*PhCry2* plasmids, and for designing gRNas for mutagenesis. We are grateful to Dr. Nipam Patel for providing access to his injection setup and to storage space for mutagenized colonies. We also thank Lauren North for help with animal husbandry and genotyping. This work was supported by grants from the National Science Foundation (NSF 2130765) to P.E and J.J.C.R and from the National Institute of General Medical Sciences (R35 GM145253) to P.E.

## Author Contributions

Conceptualization: V.L., P.E. and J.J.C.R; Methodology: V.L., Z. B., A.H., J.J.C.R. and P.E.; Investigation: V.L., Z. B., A.H.; Writing – original draft: V.L.; Writing – review & editing: V.L., P.E. and J.J.C.R, Funding acquisition: P.E. and J.J.C.R; Supervision: P.E and J.J.C.R..

## Supplemental information

**Figure S1:**
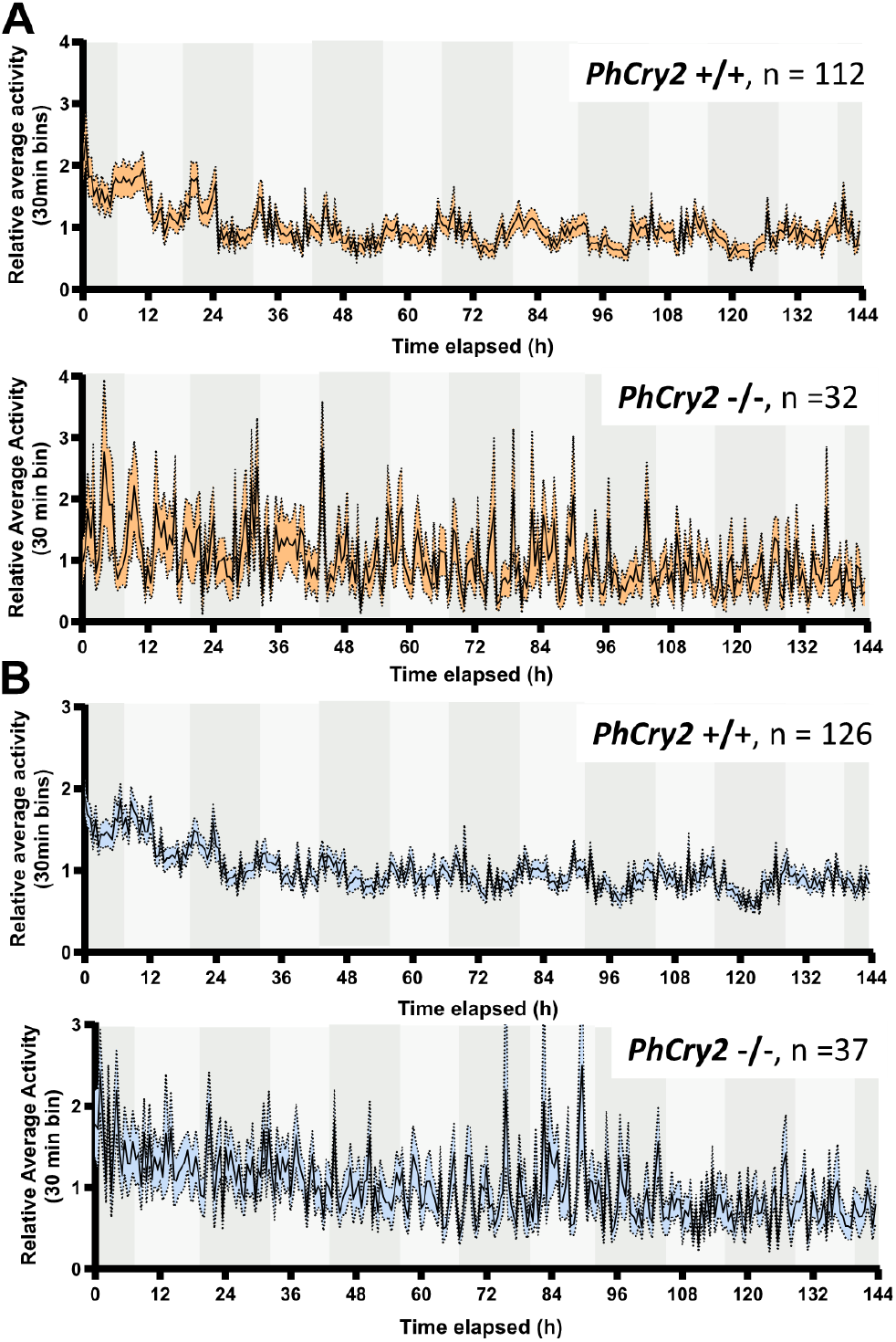
Behavior of control and *PhCry2* knock-out animals recorded in constant darkness after entrainment to LD cycles (related to Figure 1) A) Average relative vertical swimming activity of wild-type (*PhCry2*+/+) and homozygous mutant (*PhCry2*−/−) animals recorded in constant darkness (DD) after entrainment to a LD cycle. Light gray shading represents the subjective day and dark gray shading represents the subjective night. Standard error to the mean (SEM) is represented with the orange shading. B) Average relative roaming activity of wild-type (*PhCry2*+/+) and homozygous (*PhCry2*- /-) animals recorded in DD after entrainment to a LD cycle. Standard error to the mean (SEM) is represented with the blue shading.

**Figure S2:**
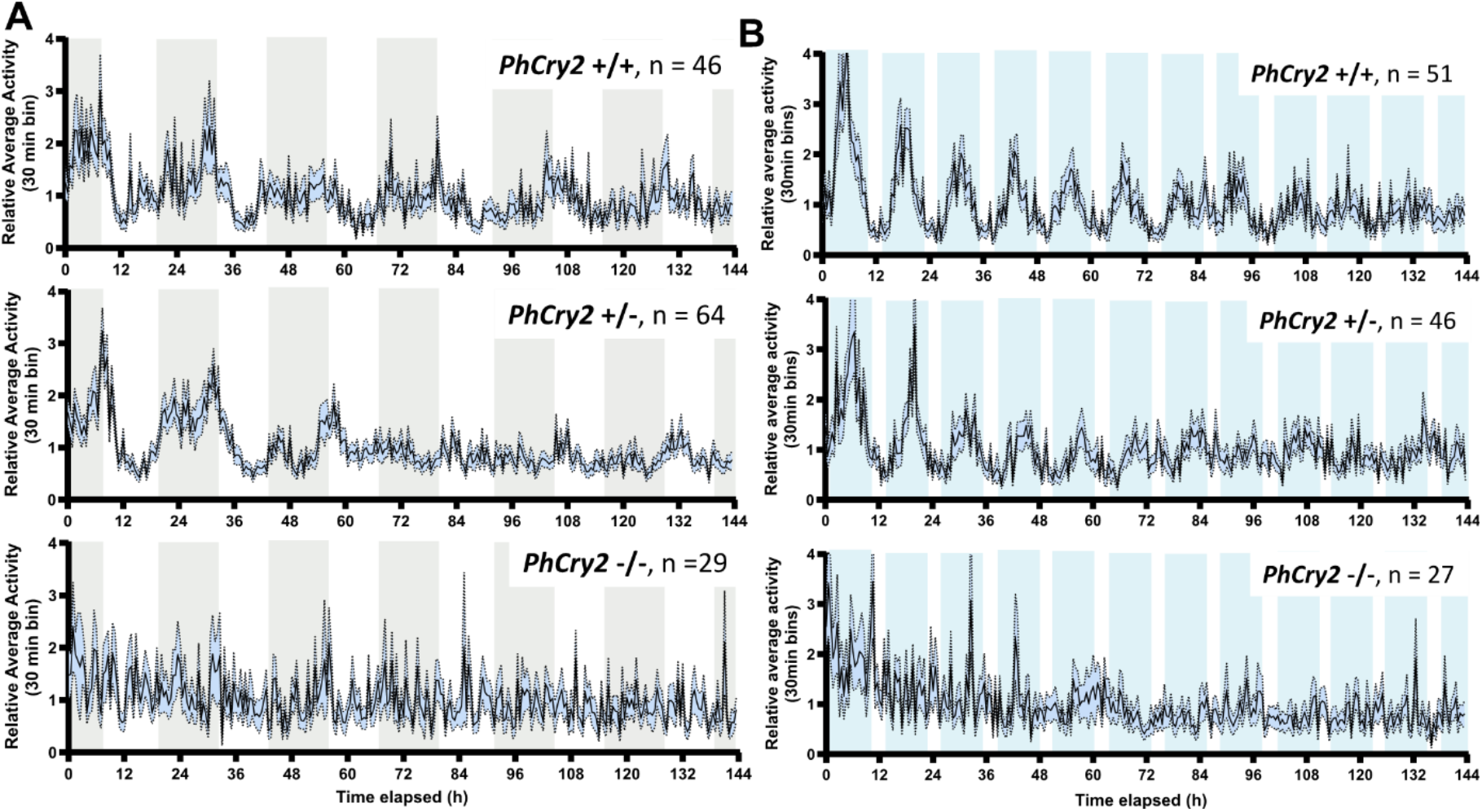
Roaming activity of *PhCry2* knock-out and control animals after LD-only or LD-tidal entrainment (Related to Figures 1 and 2) A) Average relative roaming activity of wild-type (*PhCry2*+/+), heterozygous (*PhCry2*+/-) and homozygous (*PhCry2*−/−) animals recorded in constant light (LL) after entrainment to a LD cycle. Gray shading represents the subjective night. Standard error to the mean (SEM) is represented with the blue shading. B) Average relative roaming activity of wild-type (*PhCry2*+/+), heterozygous (*PhCry2*+/-) and homozygous (*PhCry2*−/−) animals recorded in constant dark (DD) and high tide after entrainment to LD and tidal (10.3 h high tide: 2.1 h low tide) cycles. Blue shading represents the expected high tide. Standard error to the mean (SEM) is represented with the blue shading.

**Figure S3:**
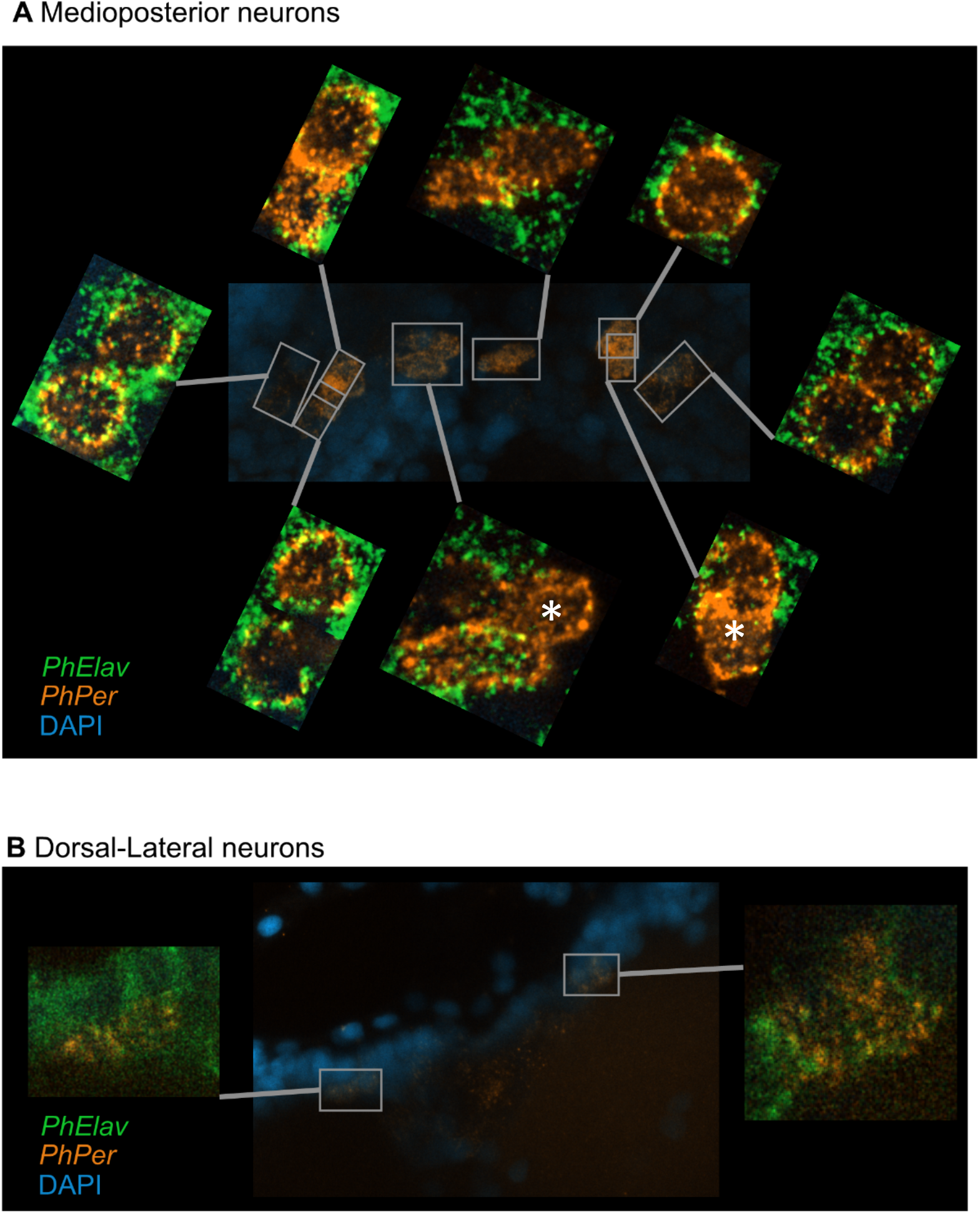
*PhPer* and *PhElav* HCR-FISH in Medioposterior and Dorsal-lateral neurons. A) Co-staining was observed in most Medioposterior cells, which are thus neurons. Two of them, however, appeared to be Elav negative and might thus be glial cells (*). N= 3 brains B) All four dorsal-lateral cells showed co-staining and are thus neurons.

**Figure S4:**
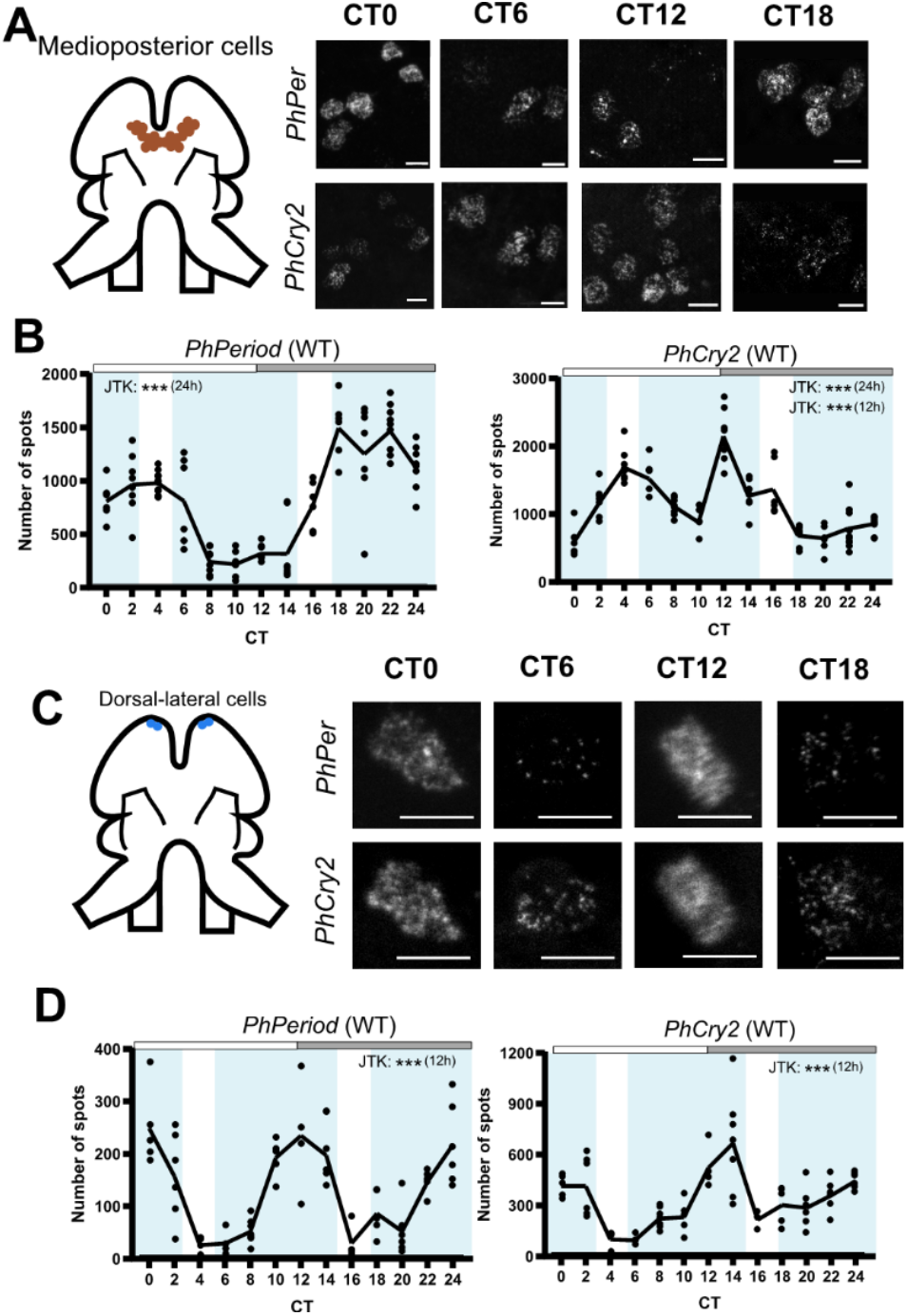
*PhPer* and *PhCry2* expression in circadian and circatidal neurons of wild type animals. A) HCR-FISH of *PhPer* and *PhCry2* mRNAs in circadian medioposterior clock neurons exposed to constant conditions after entrainment to 10.3:2.1 tidal and 12:12 LD cycles in wild type (WT) animals. A z-stack for each mRNA visualized in the left hemisphere is shown across circadian time (CT). Scale bars represent 10µm. B) Quantification of the number of HCR-FISH spots per hemisphere across CT in WT animals. Each dot represents one hemisphere of one individual and the solid line the mean. Grey shading represents the subjective dark phase and blue shading the subjective high tide. After testing the effect of time on gene expression (*PhPer*, Kruskal-Wallis, ***p <0.001, *PhCry2*, Kruskal-Wallis, ***p <0.001), rhythm was tested using the JTK-cycle algorithm for 24 h and 12 h *PhPer* mRNA rhythms (See Table S2 for exact p-value). C) HCR-FISH of *PhPer* and *PhCry2* mRNAs in circatidal dorsal-lateral clock neurons exposed to constant conditions after entrainment to 10.3:2.1 tides and 12:12 LD in WT animals as represented in A). D) As represented in B) quantification of the number of HCR-FISH spots per hemisphere across CT in WT animals. Each dot represents one hemisphere of one individual and the solid line the mean. Grey shading represents the subjective dark phase and blue shading the subjective high tide. After testing the effect of time on the gene expression (*PhPer*, Kruskal-Wallis, ***p <0.001, *PhCry2*, Kruskal-Wallis, ***p <0.001), rhythm was tested using the JTK algorithm for 24 h and 12 h (See Table S4 for exact p-value).

**Figure S5:**
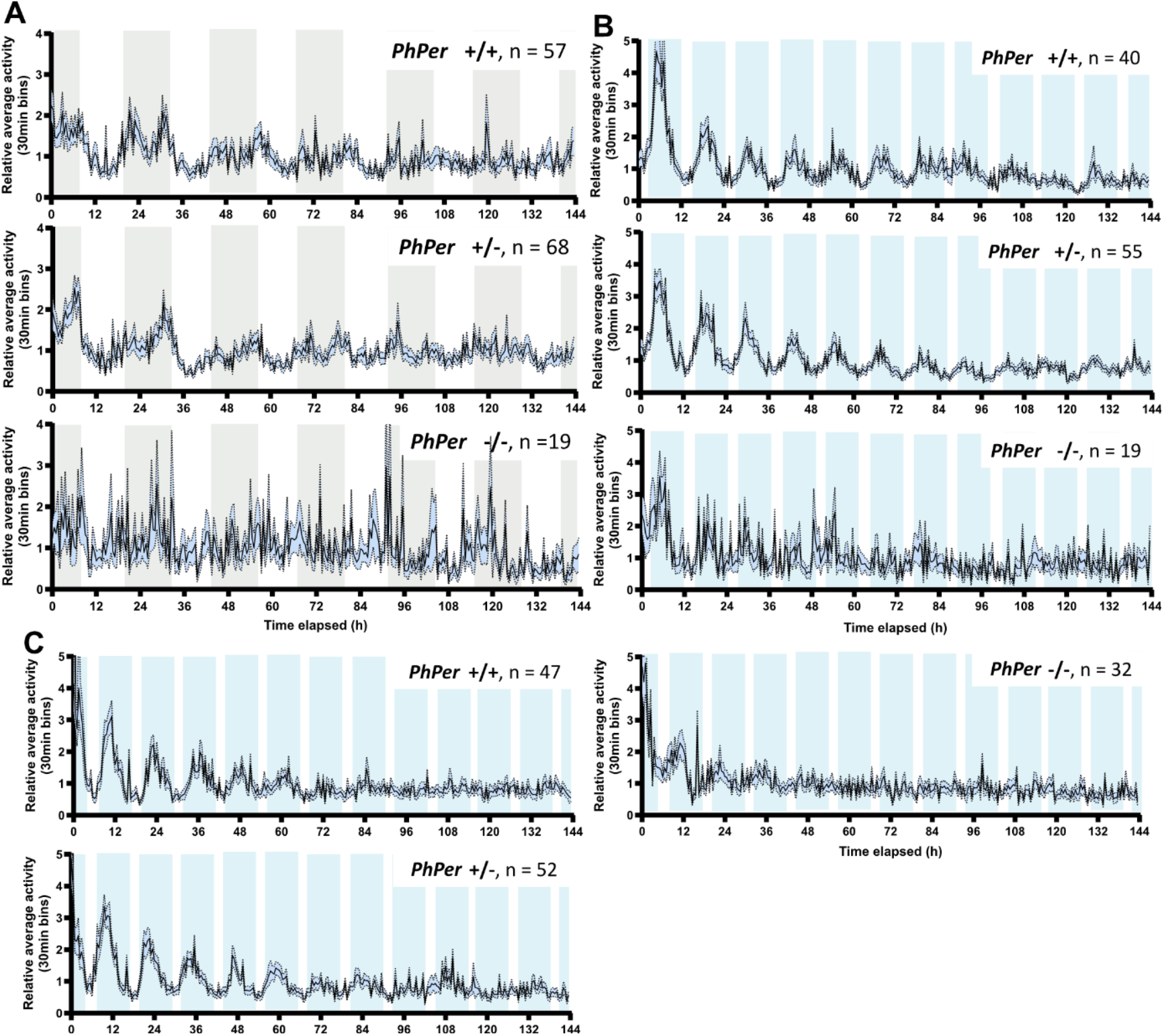
Roaming activity of *PhPer* knock-out and control animals after LD-only or LD-tidal entrainment. (Related to Figures 3 and 4) A) Average relative roaming activity of wild-type (*PhPer*+/+), heterozygous (*PhPer*+/-) and homozygous (*PhPer*−/−) animals recorded in constant light (LL) after entrainment to a LD cycle. Gray shading represents the subjective night. Standard error to the mean (SEM) is represented with the blue shading. B) Average relative roaming activity of wild-type (*PhPer*+/+), heterozygous (*PhPer*+/-) and homozygous (*PhPer*−/−) animals recorded in constant dark (DD) and high tide after entrainment to LD and tidal (10.3 h high tide: 2.1 h low tide) cycles. Blue shading represents the expected high tide. Standard error to the mean (SEM) is represented with the blue shading. The last high tide (LHT) was experienced at 13:20. C) Average relative roaming activity of wild-type (*PhPer*+/+), heterozygous (*PhPer*+/-) and homozygous (*PhPer*−/−) recorded in DD and high tide after entrainment to LD and tidal (10.3 h high tide: 2.1 h low tide) cycles as represented in B). The last high tide (LHT) was experienced at 19:20.

**Figure S6:**
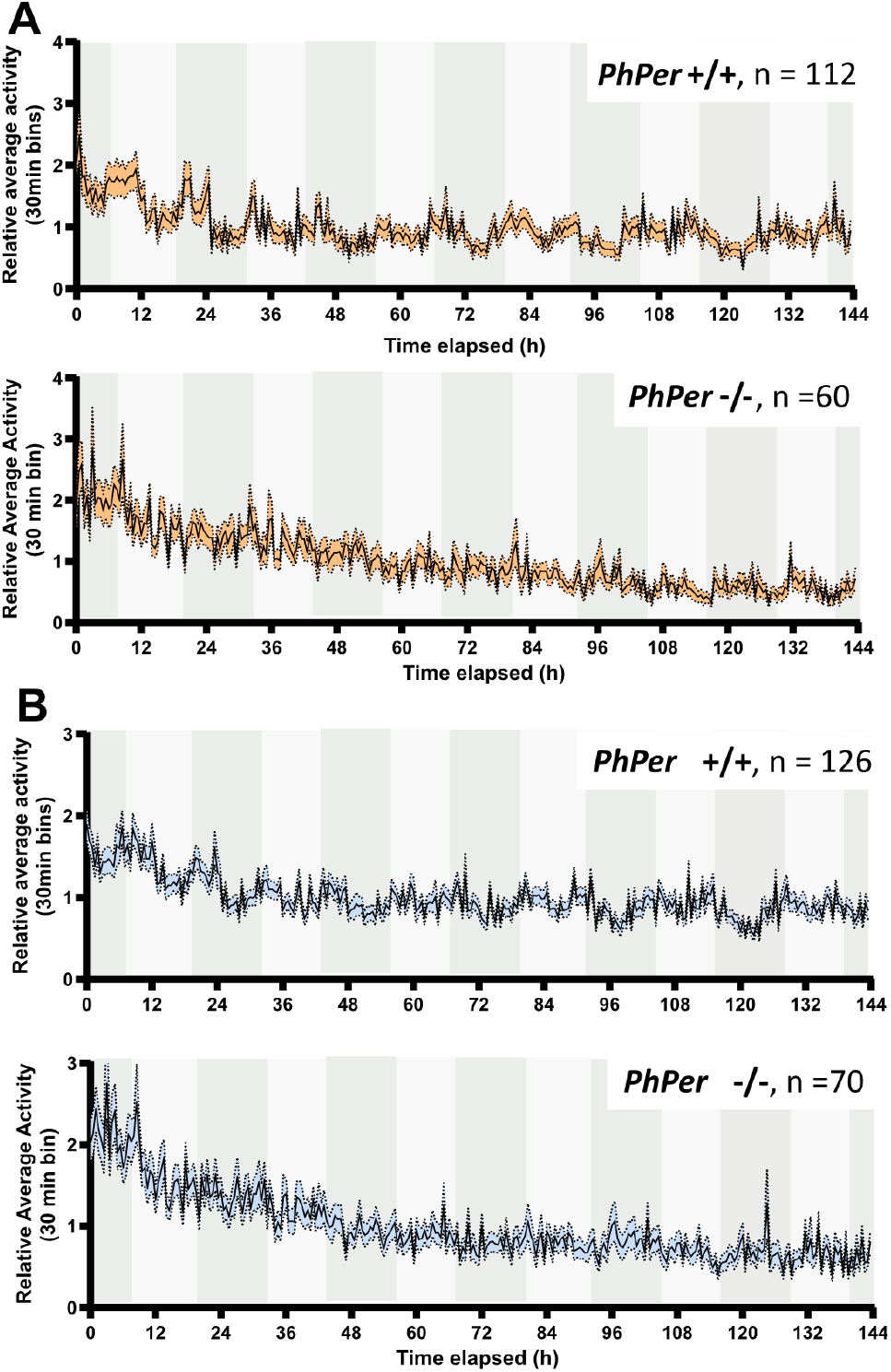
Behavior of *PhPer* knock-out and control animals recorded in constant darkness after entrainment to LD cycles (Related to Figure 3) A) Average relative vertical swimming activity of wild-type (*PhPer*+/+) and homozygous (*PhPer*−/−) animals recorded in constant darkness (DD) after entrainment to a LD cycle. Light gray shading represents the subjective day and dark gray shading represents the subjective night. Standard error to the mean (SEM) is represented with the orange shading. A) Average relative roaming activity of wild-type (*PhPer*+/+) and homozygous (*PhPer*−/−) animals recorded in DD after entrainment to a LD cycle. Standard error to the mean (SEM) is represented with the blue shading.

### Putative PAS-A Exon 1

AATTTAGATATTCCTAATGACTGTCTTCACCTCATCCCCCAGGGACTGGATGGGTTCATGCTGGCGCTCACCCTCAGCGGGGAGGCCCTCTACGTGTCGGAGAGCGTGACGTCACTGCTGGGGCACCTGCCGGTATGTACTCAACAGGCCTGGCCATCAAATGTTAAGGTAAAGTCTACTGCGAAACCCAAACGCTCTGGTTCGTGCTGCGGGAATAGGCTAACTGGTCCCAGGAAACTGTA

### Putative PAS-A Exon 2

AGAGAGAAGTAACTGGAAGGAGGTTAACGGACTTCATGCGAAAAGAAGATGGTGAAACGCTGCTCTCTAGTCTGCAAAATAGCGCAAAGAACGTCACAAGTTCTTCGAATTCAGTTTCACCACGTCTAATGACATCTACGAAAACGTCTAATAACTCTACTAGCTTAGGATTCAACCCAGCGTTCAACAACCGCGGCAATCTAAGACAAAGCGAGTTGGAGTCCACCCAGAACTTTGACCCAAGTCCTGGTGAGGTGATCCCCGTCCCAGATTTTACAGGAGGACTAGTCCCCATTTGCATCGACAGTCAGGAAAGCGTACTGCAAAGTTTTGGCGATTGCGAAAATGAATCTGTTCTTGATATTTTGCAAACAAACCAAGCACACCTGGACACGAATTCTTCTGATGTAACGTTCAATGTCCTGGATCCAACGGAAATTGGTCCAAACGGGTCCTTCCTCGAGTTCAACTCTCGACCGTTGTCAACGTTCACAGAGTCGCCAATGAAGACCCAGCAGCATCCCTCCGGCAGGGATACCAGGCCCATGTCTGCTCCTGGCTATACCAATAAATGCTATTCGGGGAAAAATGGACCGATTGGAACAAGTTCTTCGAACTGGATTAACGGTATCCAAGGATGGCCCAAGATGTCAGGTGCAAGGAAGAACACCGTTTCACCCGCTGCAGACAACACTTCTGGGTTCGGGTCAGCAAGTTCATTTTATGGAAGGGAAATATTCTATTCCGGTGCACCCATGCAAGTACAAACAGACACTACAATTTCTTCAGGATCGGATCCTGTAAATATTTGTGCTGATGACAACAAAATGGTGGAGAAATTCCCAATGGATGCCAACATGGAACATAAAAAAGGTAAGATGATAGACGTACGTCATCTTGTATTGCGATGGGAGTTTTTACGCAGTTCACTTTCTAGTTTAAACCTTGTCAACAAGAACATGAACCCTGATCGCAAAAGCTTGTAA

### Predicted PAS-A

GLDGFMLALTLSGEALYVSESVTSLLGHLPREVTGRRLTDFMRKED GETLLSSLQNSAKN

**Figure S7:**
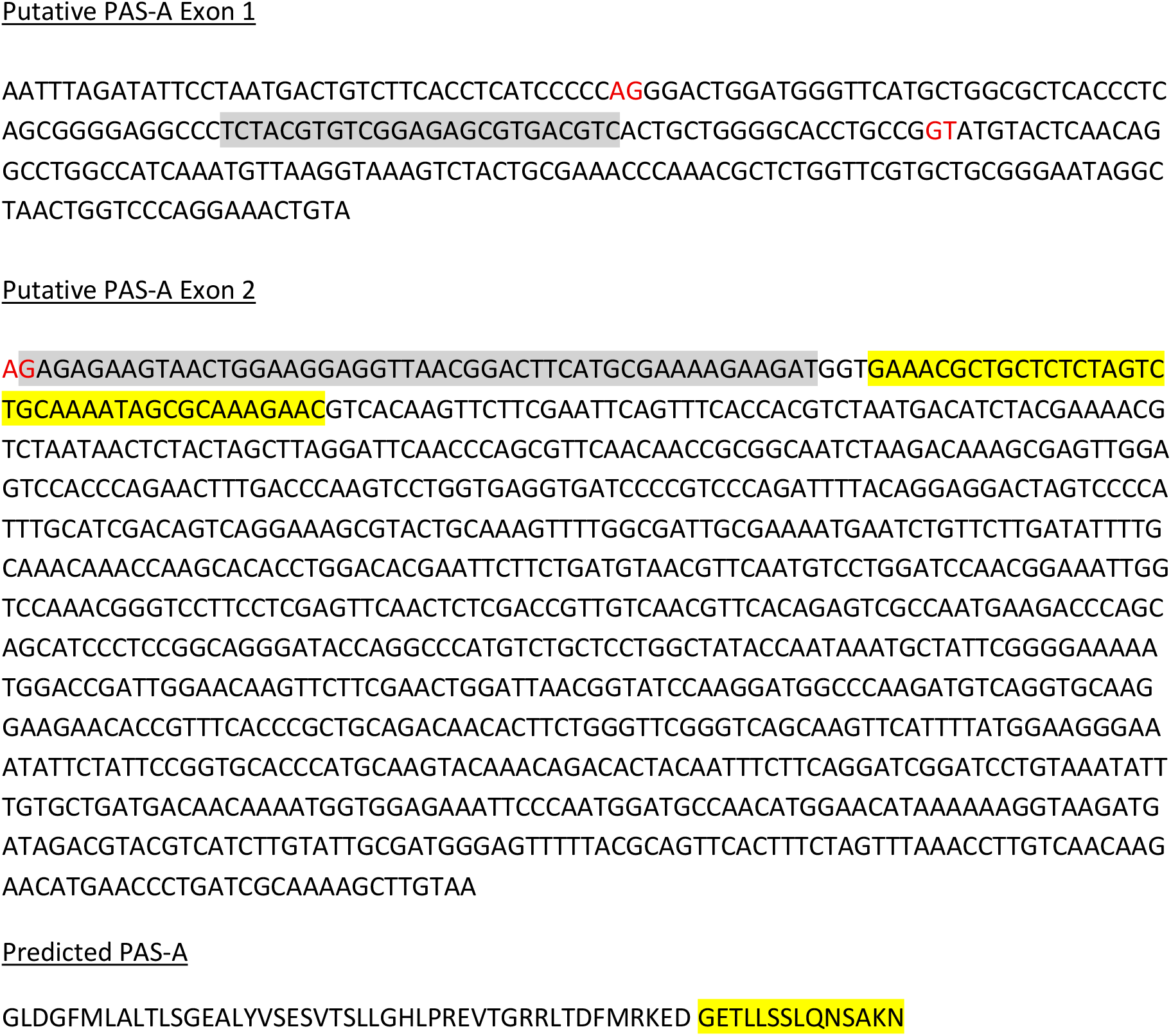
Putative *PhClk* PAS-A domain (Related to Figure 5) Two putative exons that would encode the PAS-A domain of PhCLK were identified through sequence homology. In red, predicted splicing donor and acceptor sites. The donor site of the second exon cannot be predicted in the absence of transcriptomics information, because the spacer region separating the PAS-A and PAS-B domain is poorly conserved. In grey, short transcript sequences present in the *P. hawaiensis* sequence read archive (NCBI, Exon 1: SRA:SRR17898727.161596.1, Exon 2: SRA:SRR14908235.12534108.1 and SRA:SRR060813.29331.2). In yellow, sequence showing weak homology to the region separating the PAS-A and PAS-B domain in other species.

**Figure S8:**
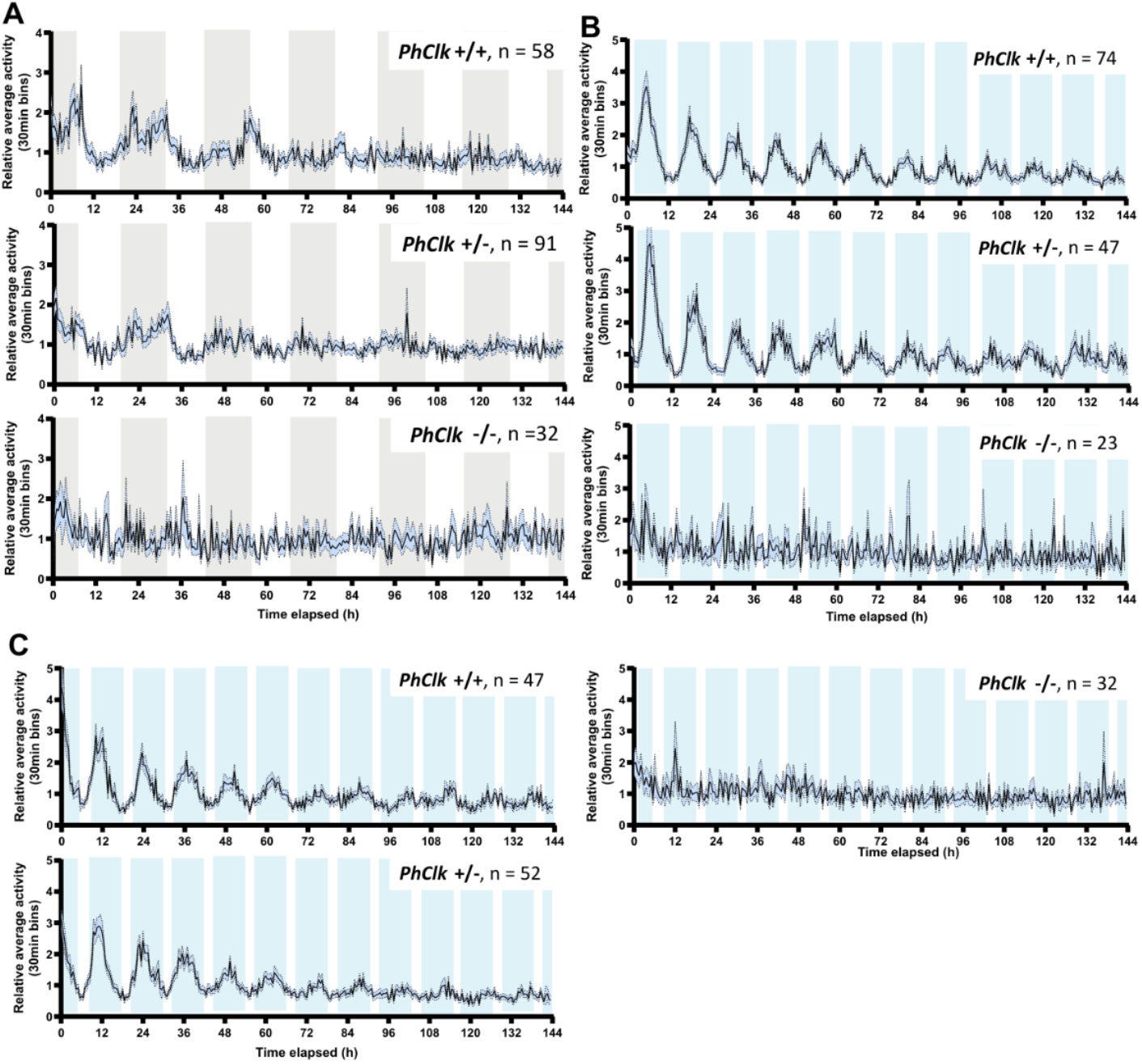
Roaming activity of *PhClk* knock-out and control animals after LD-only or LD-tidal entrainment (Related to Figures 5 and 6) A) Average relative roaming activity of wild-type (*PhClk*+/+), heterozygous (*PhClk*+/-) and homozygous (*PhClk*−/−) animals recorded in constant light (LL) after entrainment to a LD cycle. Gray shading represents the subjective night. Standard error to the mean (SEM) is represented with the blue shading. B) Average relative roaming activity of wild-type (*PhClk*+/+), heterozygous (*PhClk*+/-) and homozygous (*PhClk*−/−) animals recorded in DD and high tide after entrainment to a LD cycle and tidal (10.3 h high tide: 2.1 h low tide) cycles. Blue shading represents the expected high tide. Standard error to the mean (SEM) is represented with the blue shading. The last high tide (LHT) was experienced at 13:20. C) Average relative roaming activity of wild-type (*PhClk*+/+), heterozygous (*PhClk*+/-) and homozygous (*PhClk*−/−) animals recorded in DD and high tide after entrainment to a LD cycle and tidal (10.3 h high tide: 2.1 h low tide) cycles as represented in B). The last high tide (LHT) was experienced at 19:20.

**Figure S9:**
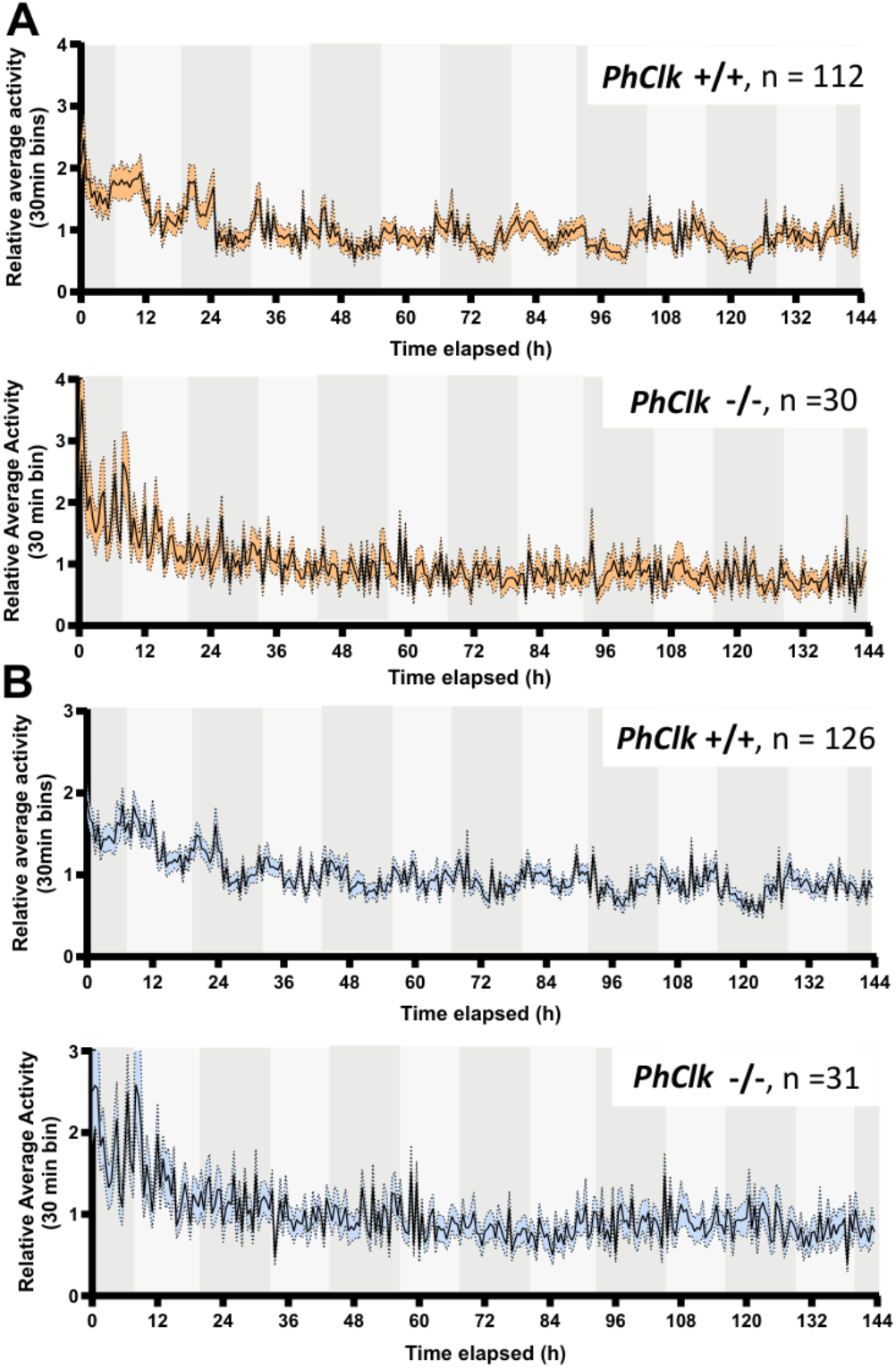
Behavior of *PhClk* knock-out and wild-type animals recorded in constant darkness after entrainment to LD cycles (Related to Figure 5) A) Average relative swimming activity wild-type (*PhClk*+/+) and homozygous (*PhClk*−/−) animals recorded in constant darkness (DD) after entrainment to a LD cycle. Light gray shading represents the subjective day and dark gray shading represents the subjective night. Standard error to the mean (SEM) is represented with the orange shading. B) Average relative roaming activity wild-type (*PhClk*+/+) and homozygous (*PhClk*−/−) animals recorded in DD after entrainment to a LD cycle. Standard error to the mean (SEM) is represented with the blue shading.

**Figure S10:**
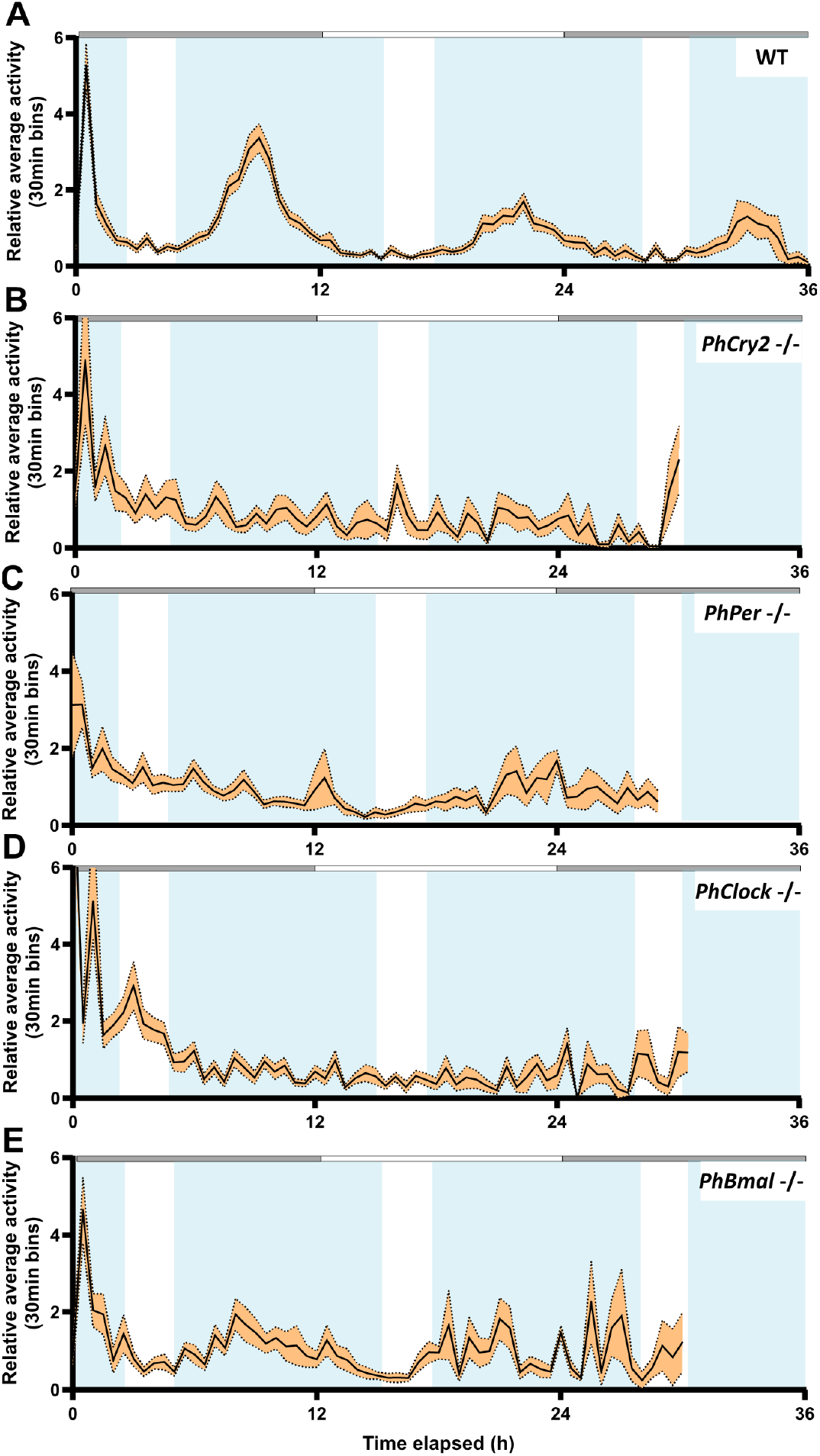
Swimming behavior of animals sampled for HCR-FISH brain staining. A) Swimming behavior of wild type (WT) animals. Grey shading represents the expected dark phase and blue shading the expected high tide. B) As represented in A), swimming behavior of *PhCry2*−/− animals. C) As represented in A), swimming behavior of *PhPer*−/− animals. D) As represented in A), swimming behavior of *PhClk*−/− animals. E) As represented in A), swimming behavior of *PhBmal1*−/− animals.

**Table S1:**
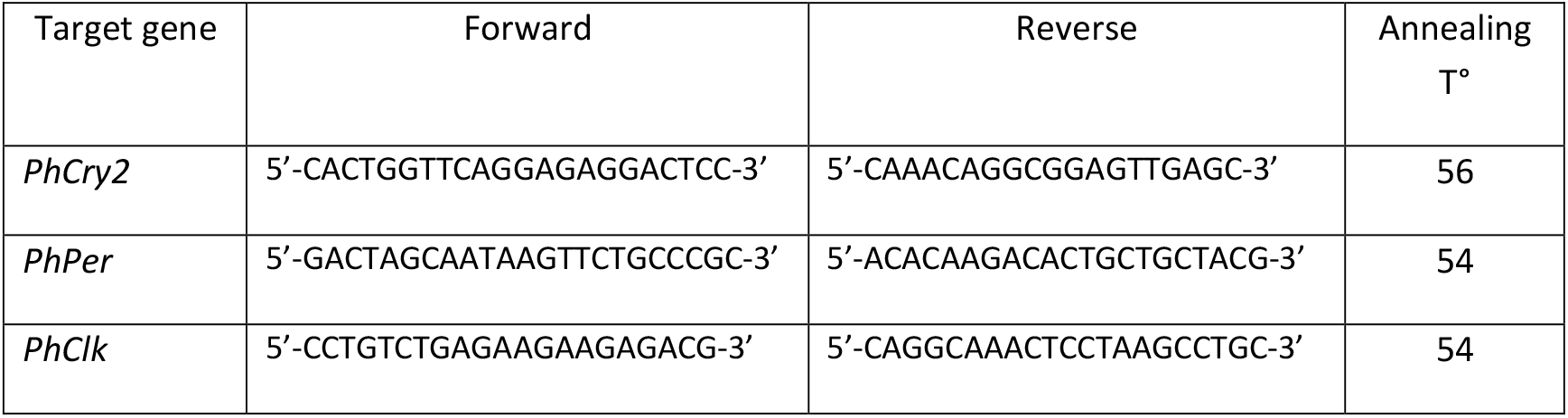
PCR primers sequences used in this study for genotyping.

**Table S2:** JTK-cycle outputs testing 24-h mRNA rhythms in Medioposterior neurons (Related to Figures 1, 3, 5 and 7)

| Genotype | Target gene | Variation through time | p-value (Bonferroni corrected) | Phase | Amplitude |
| --- | --- | --- | --- | --- | --- |
| WT<br>(13 sampling points) | *PhPer* | KW ***p<0.001 | 3.668e-17 | 22 | 484.72 |
|  | *PhCry2* | KW ***p<0.001 | 5.0561e-9 | 9 | 2411.54 |
| WT<br>(4 sampling points) | *PhPer* | ANOVA***p<0.001 | 0.00823 | 0 | 470.226 |
|  | *PhCry2* | ANOVA***p<0.001 | 1.112e-7 | 12 | 876.105 |
| *PhCry2* -/- | *PhPer* | KW **p<0.007 | 0.003698 | 3 | 222.0315 |
|  | *PhCry2* | ANOVA p = 0.205 |  |  |  |
| *PhPer* -/- | *PhPer* | KW ***p<0.001 | 0.00447 | 3 | 236.174 |
|  | *PhCry2* | ANOVA***p<0.001 | 0.03611 | 3 | 275.77 |
| *PhClk* -/- | *PhPer* | ANOVA*p = 0.013 | 0.995 | 21 | 63.639 |
|  | *PhCry2* | KW p = 0.528 |  |  |  |
| *PhBmal1* -/- | *PhPer* | ANOVA***p<0.001 | 0.0885 | 3 | 244.305 |
|  | *PhCry2* | ANOVA**p = 0.002 | 0.00374 | 12 | 140.714 |

**Table S3:**
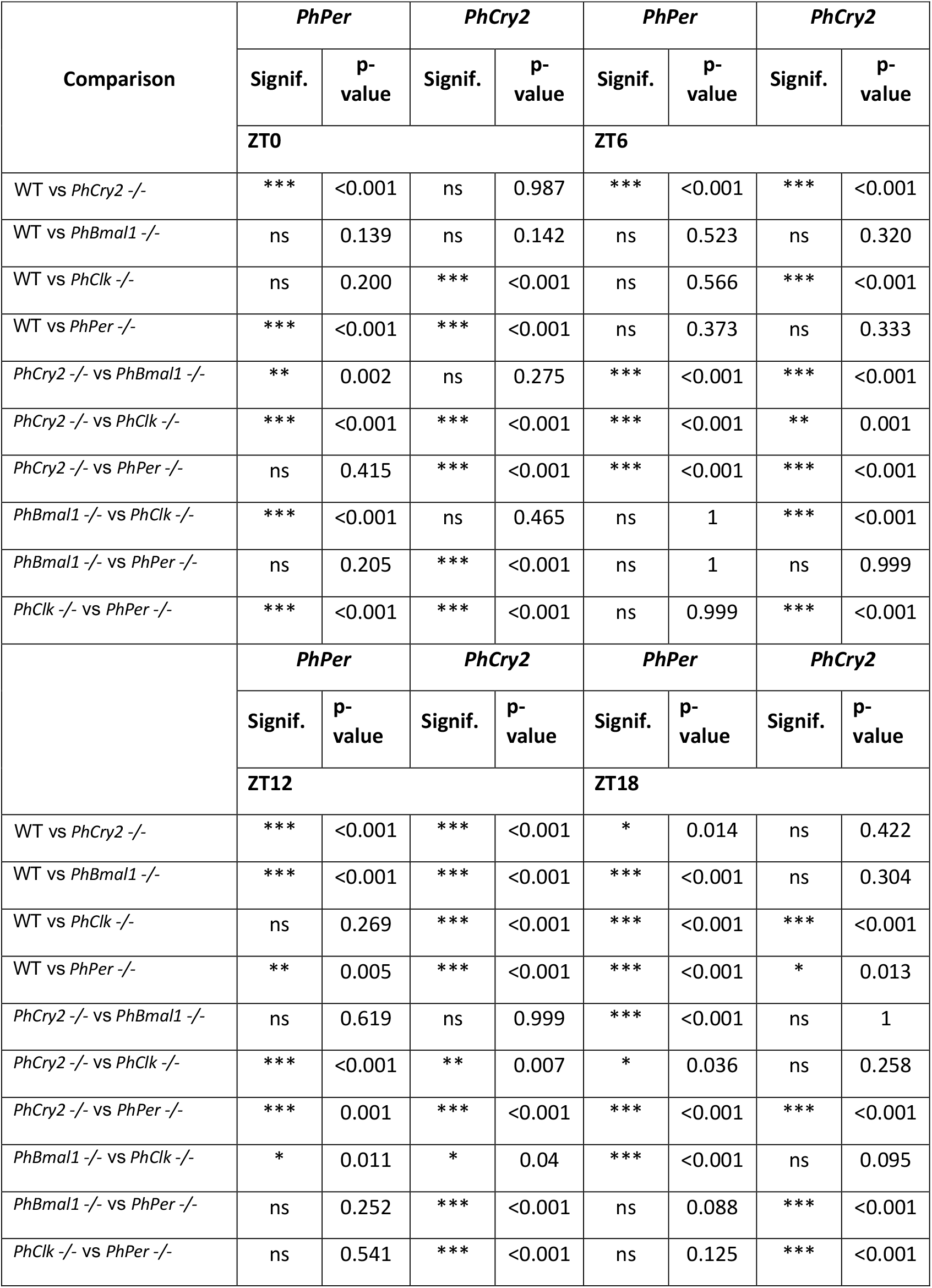
Comparison of *PhPer* and *PhCry2* mRNA levels in Medioposterior neurons as a function of genotype (Tukey’s multiple comparisons test) (Related to Figures 1, 3, 5 and 7)

**Table S4:**
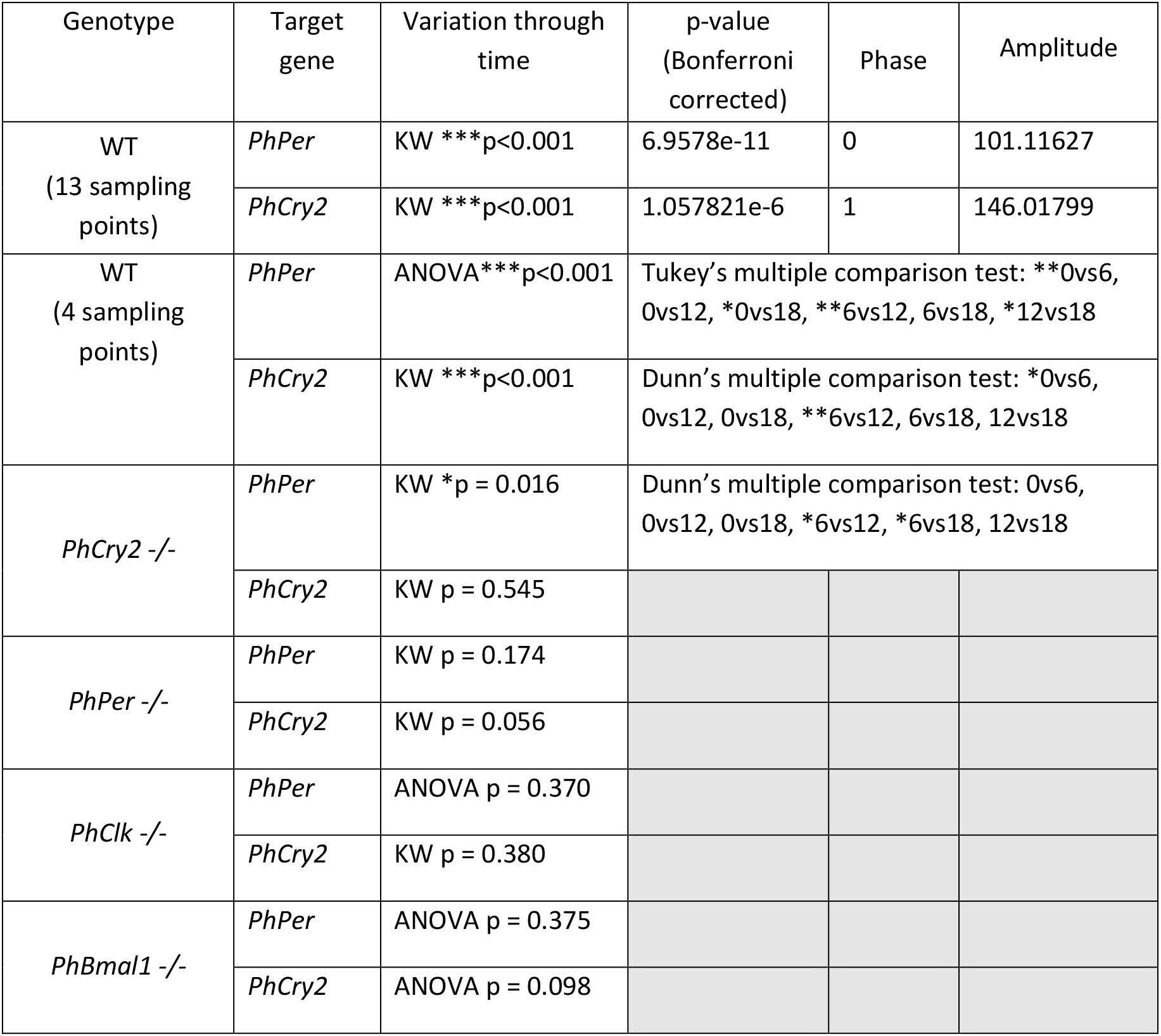
Statistical analysis testing 12-h mRNA rhythms in Dorsal-lateral neurons (Related to Figures 2, 4, 6 and 7)

| Genotype | Target gene | Variation through time | p-value (Bonferroni corrected) | Phase | Amplitude |
| --- | --- | --- | --- | --- | --- |
| WT<br>(13 sampling points) | *PhPer* | KW ***p<0.001 | 6.9578e-11 | 0 | 101.11627 |
|  | *PhCry2* | KW ***p<0.001 | 1.057821e-6 | 1 | 146.01799 |
| WT<br>(4 sampling points) | *PhPer* | ANOVA ***p<0.001 | Tukey's multiple comparison test: **0vs6, 0vs12, *0vs18, **6vs12, 6vs18, *12vs18 |  |  |
|  | *PhCry2* | KW ***p<0.001 | Dunn's multiple comparison test: *0vs6, 0vs12, 0vs18, **6vs12, 6vs18, 12vs18 |  |  |
| *PhCry2* -/- | *PhPer* | KW *p = 0.016 | Dunn's multiple comparison test: 0vs6, 0vs12, 0vs18, *6vs12, *6vs18, 12vs18 |  |  |
|  | *PhCry2* | KW p = 0.545 |  |  |  |
| *PhPer* -/- | *PhPer* | KW p = 0.174 |  |  |  |
|  | *PhCry2* | KW p = 0.056 |  |  |  |
| *PhClk* -/- | *PhPer* | ANOVA p = 0.370 |  |  |  |
|  | *PhCry2* | KW p = 0.380 |  |  |  |
| *PhBmal1* -/- | *PhPer* | ANOVA p = 0.375 |  |  |  |
|  | *PhCry2* | ANOVA p = 0.098 |  |  |  |

**Table S5:** Comparison of *PhPer* and *PhCry2* mRNA levels in Dorsal-lateral neurons as a function of genotype (Tukey’s multiple comparisons test) (Related to Figures 2, 4, 6 and 7).

| Comparison | *PhPer* |  | *PhCry2* |  | *PhPer* |  | *PhCry2* |  |
| --- | --- | --- | --- | --- | --- | --- | --- | --- |
|  | Signif. | p-value | Signif. | p-value | Signif. | p-value | Signif. | p-value |
|  | ZT0 |  |  |  | ZT6 |  |  |  |
| WT vs *PhCry2* -/- | ns | 0.091 | *** | <0.001 | *** | <0.001 | ns | 0.148 |
| WT vs *PhBmal1* -/- | *** | <0.001 | ns | 0.849 | ns | 0.097 | ns | 0.593 |
| WT vs *PhClk* -/- | ns | 1 | *** | <0.001 | *** | <0.001 | ns | 0.679 |
| WT vs *PhPer* -/- | ** | 0.005 | ns | 0.933 | ns | 0.737 | * | 0.019 |
| *PhCry2* -/- vs *PhBmal1* -/- | ns | 0.256 | *** | <0.001 | ns | 0.270 | ns | 0.982 |
| *PhCry2* -/- vs *PhClk* -/- | ns | 0.068 | ns | 0.927 | ns | 0.999 | ns | 0.796 |
| *PhCry2* -/- vs *PhPer* -/- | ns | 0.560 | ** | 0.008 | ** | <0.001 | ns | 0.914 |
| *PhBmal1* -/- vs *PhClk* -/- | *** | <0.001 | *** | <0.001 | ns | 0.175 | ns | 0.996 |
| *PhBmal1* -/- vs *PhPer* -/- | ns | 0.999 | ns | 1 | ns | 0.459 | ns | 0.724 |
| *PhClk* -/- vs *PhPer* -/- | ** | 0.004 | *** | <0.001 | *** | <0.001 | ns | 0.269 |
|  | *PhPer* |  | *PhCry2* |  | *PhPer* |  | *PhCry2* |  |
|  | Signif. | p-value | Signif. | p-value | Signif. | p-value | Signif. | p-value |
|  | ZT12 |  |  |  | ZT18 |  |  |  |
| WT vs *PhCry2* -/- | ** | 0.009 | *** | <0.001 | ns | 0.928 | ** | 0.006 |
| WT vs *PhBmal1* -/- | ns | 0.259 | *** | <0.001 | ns | 0.321 | ns | 0.467 |
| WT vs *PhClk* -/- | ns | 0.677 | *** | <0.001 | * | 0.042 | ns | 0.202 |
| WT vs *PhPer* -/- | *** | <0.001 | * | 0.022 | ns | 0.941 | ns | 0.886 |
| *PhCry2* -/- vs *PhBmal1* -/- | ns | 0.935 | ns | 0.188 | ns | 0.804 | ns | 0.344 |
| *PhCry2* -/- vs *PhClk* -/- | ns | 0.231 | ns | 0.921 | ns | 0.251 | ns | 0.658 |
| *PhCry2* -/- vs *PhPer* -/- | ns | 0.867 | *** | <0.001 | ns | 0.523 | ns | 0.080 |
| *PhBmal1* -/- vs *PhClk* -/- | ns | 0.892 | ns | 0.6332 | ns | 0.877 | ns | 0.986 |
| *PhBmal1* -/- vs *PhPer* -/- | ns | 0.567 | ns | 0.248 | ns | 0.067 | ns | 0.949 |
| *PhClk* -/- vs *PhPer* -/- | * | 0.037 | *** | <0.001 | ** | 0.005 | ns | 0.727 |

**Table S6:** Period and power of locomotor behavior. Mean and SEM are indicated. (Related to Figures 1, 2, 3, 4, 5 and 6)

| Genotype | Power | Period doublet (h) | Period singlet (h) |
| --- | --- | --- | --- |
| Entrainment to LD cycles recorded in LL |  |  |  |
| *PhCry2* +/+ | 77.7 ± 9.62 | 25.4 ± 0.369 | 11.4 ± 1.40 |
| *PhCry2* +/- | 66.2 ± 6.93 | 25.2 ± 0.185 | 11.8 ± 1.50 |
| *PhCry2* -/- | 31.8 ± 6.63 | 27.4 ± 2.46 | 12.3 ± 4.30 |
| *PhPer* +/+ | 53.9 ± 9.71 | 25.6 ± 0.480 | 12.3 ± 0.051 |
| *PhPer* +/- | 57.5 ± 6.36 | 24.7 ± 0.136 | 12.3 ± 0.102 |
| *PhPer* -/- | 16 ± 4.3 | 25.6 ± 1.10 | 12.4 ± 0 |
| *PhClk* +/+ | 60.7 ± 7.46 | 25.0 ± 0.233 | 11.7 ± 1.1 |
| *PhClk* +/- | 55.8 ± 8.27 | 24.9 ± 0.674 | 12.2 ± 0.9 |
| *PhClk* -/- | 31.6 ± 6.43 | 27.6 ± 2.61 | 11.9 ± 3 |
| *PhCry2* +/+ | 71.2 ± 8.33 | 24.8 ± 0.142 | 12.4 ± 0.101 |
| Entrainment to LD cycles and tides (LHT 13 h20) recorded in DD and cst HT |  |  |  |
| *PhCry2* +/+ | 74 ± 6.25 | 25.1 ± 0.071 | 12.6 ± 0.036 |
| *PhCry2* +/- | 83.5 ± 9.75 | 25.4 ± 0.089 | 12.7 ± 0.042 |
| *PhCry2* -/- | ND | ND | ND |
| *PhPer* +/+ | 60.7 ± 4.91 | 25.06 ± 0.08 | 12.5 ± 0.05 |
| *PhPer* +/- | 64.0 ± 4.46 | 24.8 ± 0.06 | 12.4 ± 0.03 |
| *PhPer* -/- | 39.5 ± 7.60 | 25.5 ± 1.22 | 13.3 ± 0.97 |
| *PhClk* +/+ | 77.7 ± 7.12 | 25.0 ± 0.075 | 12.5 ± 0.032 |
| *PhClk* +/- | 80.1 ± 8.87 | 24.8 ± 0.110 | 12.4 ± 0.066 |
| *PhClk* -/- | 37.4 ± 11.12 | 23.6 ± 0.862 | 13.9 ± 1.41 |
| *PhCry2* +/+ | 58.7 ± 5.04 | 24.8 ± 0.09 | 12.5 ± 0.05 |
| Entrainment to LD cycles recorded in DD |  |  |  |
| WT | 54.4 ± 4.72 | 24.5 ± 0.09 | 12.3 ± 0.051 |
| *PhCry2* -/- | 28.9 ± 11.8 | 26.6 ± 2.60 | 12.1 ± 0 |
| *PhPer* -/- | 68.1 ± 15.7 | 23.3 ± 0.382 | 11.8 ± 0.166 |
| *PhClk* -/- | ND | ND | ND |
| Entrainment to LD cycles and tides (LHT 19 h20) recorded in DD and cst HT |  |  |  |
| *PhPer* +/+ | 64.3 ± 7.13 | 25.0 ± 0.118 | 12.5 ± 0.08 |
| *PhPer* +/- | 62.0 ± 6.45 | 25.2 ± 0.417 | 12.4 ± 0.05 |
| *PhPer* -/- | 29.3 ± 10.9 | 26.3 ± 3.081 | 13.5 ± 1.35 |
| *PhClk* +/+ | 65.85 ± 6.86 | 25.3 ± 0.143 | 12.6 ± 0.06 |
| *PhClk* +/- | 65.4 ± 10.2 | 24.9 ± 0.142 | 12.5 ± 0.06 |
| *PhClk* -/- | 11.8 ± 0 | 26.3 ± 0 | ND |

